# Flotillins affect LPS-induced TLR4 signaling by modulating the trafficking and abundance of CD14

**DOI:** 10.1101/2023.12.11.571077

**Authors:** Orest V. Matveichuk, Anna Ciesielska, Aneta Hromada-Judycka, Natalia Nowak, Ichrak Ben Amor, Katarzyna Kwiatkowska

**Author notes:** Equal contribution (Orest V. Matveichuk, Anna Ciesielska). Orest V. Matveichuk; ORCID 0000-0002-4057-7598 Anna Ciesielska; ORCID 0000-0001-9902-494X Aneta Hromada-Judycka; ORCID 0000-0002-4449-882X Natalia Nowak; ORCID 0000-0003-3140-2056 Ichrak Ben Amor; ORCID 0000-0003-3263-7164 Katarzyna Kwiatkowska; ORCID 0000-0002-0550-8394.

## Abstract

Lipopolysaccharide induces a strong pro-inflammatory reaction of macrophages upon activation of Toll-like 4 receptor (TLR4) with the assistance of CD14 protein. Considering a key role of plasma membrane rafts in CD14 and TLR4 activity and the significant impact exerted on that activity by endocytosis and intracellular trafficking of the both LPS acceptors, it seemed likely that the pro-inflammatory reaction could be modulated by flotillins. Flotillin-1 and -2 are scaffolding proteins associated with plasma membrane rafts and also with endo-membranes, affecting both the plasma membrane dynamics and intracellular protein trafficking. To verify the above hypothesis, a set of shRNA was used to down-regulate flotillin-2 in Raw264 cells, which were found to also become deficient in flotillin-1. The flotillin deficiency inhibited strongly the TRIF-dependent endosomal signaling of LPS-activated TLR4, and to a lower extent also the MyD88-dependent one, without affecting the cellular level of TLR4. In contrast, the depletion of flotillins down-regulated the CD14 mRNA level and the total cellular content of CD14 protein, and decreased the amount of CD14 on the cell surface. The constitutive CD14 endocytosis remained unchanged but CD14 recycling was enhanced via EEA1-positive early endosomes and golgin-97-positive *trans*-Golgi network, likely to compensate for the depletion of the cell-surface CD14. Notably, a paucity of surface CD14 in resting cells can inhibit TLR4 signaling after the stimulation of cells with LPS. In conclusion, we have shown here that flotillins modulate the pro-inflammatory response of macrophages to LPS by affecting the abundance of CD14.

## Introduction

Lipopolysaccharide (LPS) is a main constituent of the outer membrane of Gram-negative bacteria. Upon infection, LPS triggers pro-inflammatory signaling pathways activating Toll-like receptor 4 (TLR4) present in the plasma membrane of numerous immune cells, including macrophages, and some non-immune cells. If released to the cytosol, LPS binds to caspases 4/5/11 inducing programmed pro-inflammatory cell death called pyroptosis [1, 2]. Local acute inflammation helps to combat the infection, however, an overwhelming body’s response to infection leads to sepsis with a potentially fatal outcome. On the other hand, prolonged low-grade inflammation contributes to the development of type 2 diabetes, cardiovascular disease, and other diseases[3, 4]. Such chronic inflammation is potentiated by the activity of cytoplasmic NLRP3 inflammasome, which can be activated downstream of TLR4 signaling [5].

TLR4 activation is a multi-step process which starts from the recognition of LPS by lipopolysaccharide-binding protein (LBP) and next by the CD14 protein [6]. CD14 exists in two forms – a soluble protein (sCD14) found in serum, and a plasma membrane-bound glycosylphosphatidylinositol (GPI)-anchored protein (mCD14, CD14) present mainly in myeloid cells, including monocytes, macrophages, and dendritic cells. Both forms of CD14 bind LPS and are engaged in TLR4 activation. Notably, the involvement of the membrane-bound CD14 at this stage of TLR4 activation can be dispensable at higher LPS concentrations due to the likely contribution of albumin and sCD14 in delivering LPS to the receptor [7, 8]. Upon LPS binding, CD14 dissociates from the LPS/LBP complex and transfers LPS monomers to the MD2 protein of the TLR4/MD2 heterodimer. Thus activated, TLR4 recruits two adaptor proteins, TIRAP and MyD88, triggering a signaling pathway leading ultimately to the activation of the NFκB, AP-1 and CREB transcription factors. A hallmark pro-inflammatory cytokine expressed in this pathway is TNFα [9–11]. Also, glycolysis is induced in this MyD88-dependent way to provide cells with metabolic intermediates required for gene expression and cytokine production [12, 13]. Subsequently, the complex of TLR4/MD2 with LPS is internalized and in endosomes TLR4 recruits a second pair of adaptor proteins, TRAM and TRIF. This triggers the second signaling pathway of TLR4 which leads to the activation of the IRF3/7 transcription factors and the production of type I interferons, chemokine CCL5/RANTES, and proteins encoded by interferon-regulated genes, like IP-10. Also, so-called late-phase activation of NFκB takes place [9, 14, 15].

It has been established that TLR4/LPS endocytosis is governed by the membrane-bound CD14 and several regulators of the endocytosis have been revealed, including enzymes catalyzing the turnover of phosphatidylinositol 4,5-bisphosphate [PI(4,5)P_2_] [16–18]. We found that CD14 itself determines the synthesis of PI(4,5)P_2_ in LPS-stimulated cells [19]. Recently, a role of CD14 in the Ca^+2^ influx mediated by the plasma membrane TRPM7 channel necessary for TLR4 endocytosis in macrophages has been demonstrated [20]. Notably, also in unstimulated macrophages CD14 undergoes slow constitutive endocytosis and is degraded in lysosomes, and its *de novo* synthesis is required to replenish the cell surface pool of CD14. Upon LPS binding, the constitutive CD14 endocytosis is accelerated concomitant with TLR4/MD2/LPS uptake [21]. Additionally, our recent studies have revealed that a fraction of the endocytosed CD14 can recycle back to the plasma membrane, both in resting and in LPS-stimulated cells, via a pathway depending on sorting nexins SNX1, 2, and 6. This newly discovered pathway contributes to the maintenance of the cell surface pool of CD14 determining the extent of the activation of TLR4 by LPS [22]. This is a good illustration of how the ultimate reaction of macrophages to LPS is modulated by diverse proteins controlling the cellular trafficking of CD14. The CD14 trafficking through the endo-lysosomal route is divergent from that of TLR4, which increases the likelihood of a dysregulation of the inflammatory reactions to LPS by a malfunctioning of this compartment that is intrinsic to several human hereditary diseases [23].

The contribution of CD14 to TLR4 activation and endocytosis in macrophages is likely related to its localization in distinct nanodomains of the plasma membrane enriched in sphingolipids and cholesterol called rafts. The ample data supporting this assumption include bio-imaging assays, the sensitivity of LPS-induced TLR4 signaling to changes of cell cholesterol level, and studies of the protein content of the so-called detergent-resistant membrane (DRM) fraction roughly corresponding to rafts [24]. Also, a selected group of proteins that associate with rafts, like the Lyn tyrosine kinase, have been proposed to regulate TLR4 activity [25]. We have recently found that stimulation of Raw264 macrophage-like cells with LPS upregulates the level of *S*-palmitoylated flotillin-1 [26]. Flotillin-1 (reggie-2) and flotillin-2 (reggie-1) are widely expressed proteins of about 50 kDa often considered as raft markers. This association can be mediated by hydrophobic interactions of flotillins with membrane lipids, including sphingosine and cholesterol, and additionally facilitated by their *S*-palmitoylation [27]. Reciprocally, flotillins can contribute to the raft assembly by acting in cooperation with the scaffolding protein MPP1[28]. The membrane-binding ability is attributed to the N-terminal fragment of the SPFH domain, so named after proteins containing a homologous sequence, ca. 200 amino acids long. The following extended α-helical part of flotillins mediates their oligomerization and is also involved in numerous intermolecular interactions[27]. Due to their membrane binding on the one hand and direct binding of numerous proteins on the other, homo- and hetero-oligomers of flotillins can serve as scaffolds enabling the assembly of multiprotein complexes involved in signal transduction, clathrin-/caveolin-independent endocytosis of plasma membrane proteins, and cytoskeleton remodeling [29–34]. Importantly, a growing set of recent data indicates that flotillins can also associate with endo-membranes, including the Golgi apparatus and the endo-lysosomal compartment, to take part in protein trafficking and thereby affect signaling of diverse plasma membrane receptors [35–39].

Surprisingly, the data on the contribution of flotillins to the signaling of TLRs are scarce. Flotillins are used as markers of rafts in immunohistochemical or biochemical analyses, and their co-localization with TLR4 suggested an association of activated TLR4 with these plasma membrane domains detected in liver samples and in Kupffer cells [40]. Flotillin-1 was found to modulate TLR3 signaling in HUVEC endothelial cells by affecting caveolin-1 level [41]. A down-regulation of stomatin-like protein-2, a member of the SPFH family, in Raw264 cells was detrimental for the raft integrity, as indicated, i.a., by a depletion of the DRM fraction of flotillins, CD14, and MyD88. As a result, activation of various TLRs, including TLR4, was inhibited [42].

In this work, we analyzed the involvement of flotillins in TLR4 pro-inflammatory activity and found that they are crucial for maximal TRIF-dependent endosomal signaling of TLR4 due to their pleiotropic influence on the level, trafficking, and localization of CD14 protein.

## Materials and Methods

### Cell culture and stimulation

Raw264.7 (further Raw264) cells and J774 cells (both from ATTC) were cultured in DMEM containing 10% FBS (Thermo Fisher Scientific) with 4.5 and 1 g/l glucose, respectively. Thioglycolate-elicited macrophages were isolated from 8- to 14-week-old male C57BL/6 mice, as described previously [22]. The procedure had been reviewed and approved by the Local Animal Ethics Committee (permission No. 394/2017). For stimulation, Raw264 cells were overlaid with fresh DMEM/10% FBS supplemented with 10 or 100 ng/ml smooth LPS from *Escherichia coli* O111:B4 (List Biological Laboratories) and incubated for up to 6 h (5% CO_2_, 37°C).

### Preparation of Raw264 cells with stably silenced *Flot2*

Cells were infected separately with five different lentiviral transduction particles (at MOI = 5) containing five different shRNA species (Merck) specific for the mouse *Flot2* gene (NM_008028, Table 1). Cells infected in two independent attempts with non-mammalian shRNA transduction particles (Merck) served as controls. The cells were cultured in the presence of 2 µg/ml puromycin (Merck) until the mortality of non-infected cells reached 100% leaving cells transfected with shRNA alive. The efficiency of the *Flot2* silencing was verified with RT-qPCR using primers specific to the mouse *Flot2* gene, with *Tbp* as reference (primers listed in Table 2).

**Table 1.**
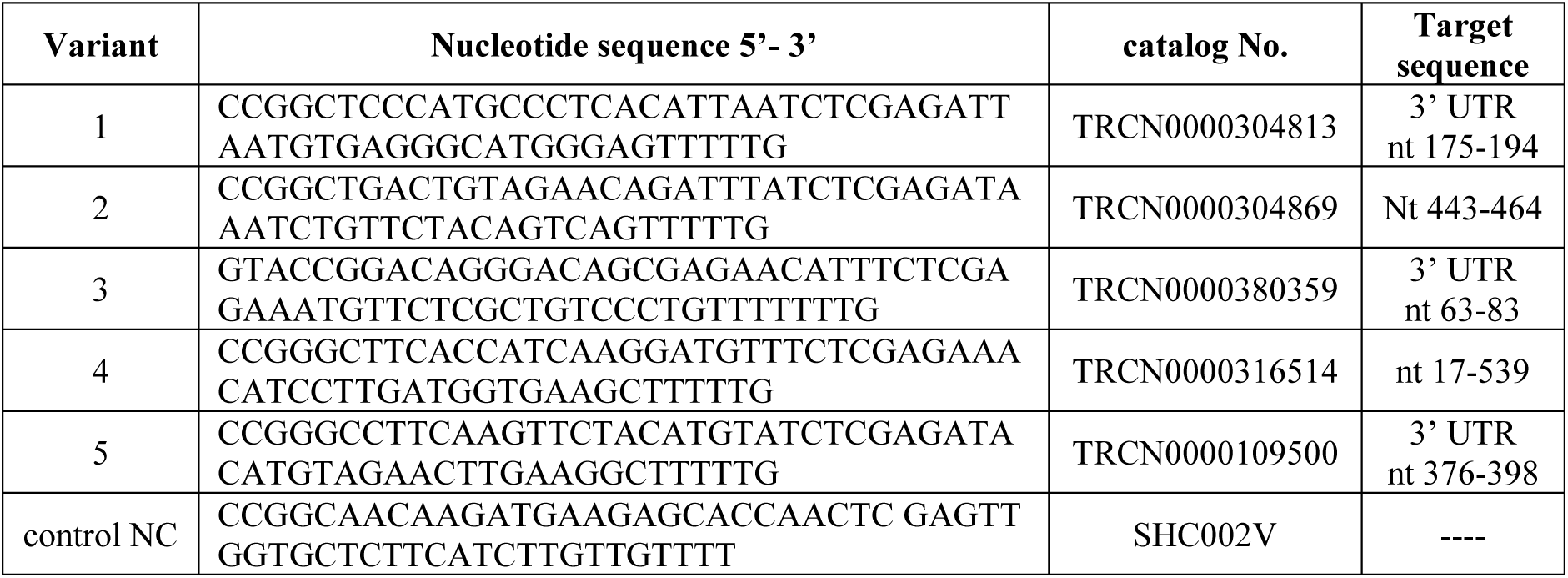
shRNA species used to silence *Flot2*.

**Table 2.**
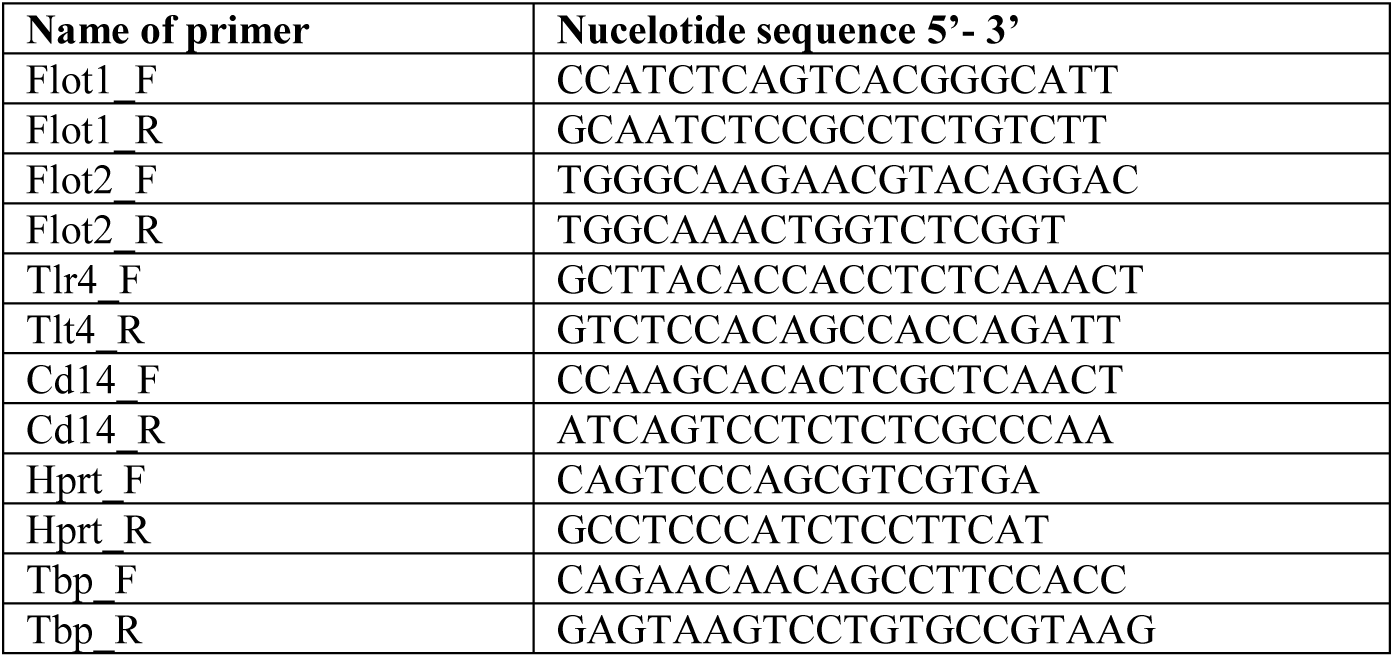
List of used primers.

### Cytokine and sCD14 assays

TNFα and CCL5/RANTES were quantified in cell culture supernatants using mouse ELISA kits (BioLegend, R&D Systems) according to manufacturers’ instructions. Samples were run in triplicates and normalized against protein content in the cells determined by crystal violet staining. sCD14 was determined in cell supernatants by mouse ELISA kit (R&D System) according to manufacturer’s instructions. The absorbance was read in a Sunrise plate reader (Tecan). Alternatively, supernatants from triplicate cultures were combined and analyzed using the Mouse Cytokine Array Kit, Panel A (R&D System). In this assay, the profile of secreted inflammatory markers was quantified by incubation with a cytokine array membrane followed by immunoblotting performed according to the manufacturer’s instructions. Immunoreactive dots were subjected to a densitometric analysis using the ImageJ program and normalized against internal standards present on the membranes.

### CD14 endocytosis

A biotinylation-based assay was used to assess CD14 endocytosis essentially as described earlier [22]. Briefly, Raw264 cells (1.5 x 10^6^ per 3.5-cm dish) were grown for 24 h, the culture medium was exchanged for a fresh one, and after 2 h the cells were washed and overlaid with ice-cold PBS containing 1 mM MgCl_2_ and 0.1 mM CaCl_2_, pH 8.2 (PBS+) supplemented with 0.5 mg/mL EZLink Sulfo-NHS-SS-Biotin (30 min on ice; Thermo Fisher Scientific). Subsequently, the excess of biotin was removed by washing the cells with PBS+ containing 5% FBS and 25 mM lysine, and twice with ice-cold PBS+. Each variant was run in duplicate and at this stage of the procedure one of the samples was lysed (30 min on ice) in 300 μl of a lysis buffer (25 mM HEPES, 0.5% SDS, 0.5% NP-40, 10% glycerol; pH 8.2) containing inhibitors (1 mM PMSF, 2 μg/ml aprotinin, 2 μg/ml leupeptin, 1 mM Na_3_VO_4_, 50 μM PAO, 10 mM *p*-nitrophenyl phosphate) and 250 units of Benzonase (Merck). An aliquot of 50 μl was withdrawn from each lysate for 10% SDS-PAGE and immunoblotting analysis of the input CD14, while the rest served to determine the level of surface-bound CD14, as described below. The other sample was incubated in a complete culture medium for 10 min at 37 °C to induce endocytosis. To remove the biotin tag remaining on the cell surface, the cells were washed with PBS+ and incubated twice (15 min each) with 50 mM MESNA (Merck) in 100 mM Tris, 100 mM NaCl, pH 8.6, followed by a wash with 5 mg/ml iodoacetamide in PBS+. Finally, the cells were washed with PBS+ and lysed (30 min on ice) in 300 μl of the lysis buffer. An aliquot of 50 μl was withdrawn from each lysate to analyze the input CD14 as above. The remaining portion of these lysates, as well as of the lysates obtained after cell surface biotinylation, was diluted 5 times in the lysis buffer without detergents and incubated with streptavidin beads (30 μl per sample, High Capacity Streptavidin Resin, Thermo Fisher Scientific) with agitation overnight at 4°C. Next, the beads were washed three times with the lysis buffer containing 0.1% SDS and 0.1% NP-40, and bound proteins were eluted by incubation for 15 min at 95 °C in SDS-sample buffer containing 2% β-mercaptoethanol, and the released proteins were subjected to 10% SDS-PAGE and immunoblotting for CD14.

### CD14 recycling assay

Raw264 cells (0.7 x 10^5^ per 15×15 mm coverslip) were grown for 24 h, next the culture medium was exchanged for a fresh one for 2 h. The assay for CD14 recycling was performed essentially as described earlier [22]. Briefly, cells were incubated with anti-mouse CD14 rat IgG2a (clone Sa14-2; BioLegend) for 30 min at 4 °C, and transferred to 37 °C for 30 min to induce CD14 uptake. Next, the cells were washed twice (2 min each) with ice-cold DMEM, pH 2.0, and twice with ice-cold PBS, pH 7.4, and incubated in the complete medium for 30 min at 37 °C to allow CD14/IgG recycling, and next for 30 min at 37 °C in the presence of chicken anti-rat IgG conjugated with Alexa Fluor 647 (Invitrogen) to stain CD14/IgG on the cell surface and allow its subsequent uptake. In control samples, anti-CD14 was replaced by the corresponding rat isotype IgG2a (BioLegend) or the antibody was omitted. The cells were then washed twice (2 min each) with ice-cold DMEM, pH 2.0, and twice with ice-cold PBS, pH 7.4, fixed with 4% formaldehyde (20 min, room temp.), permeabilized with 0.1% saponin in PBS and incubated with 4 μg/ml Hoechst 33342 (Merck) for 10 min at room temp. in PBS. To assess the initial surface level of CD14, the cells were incubated with anti-mouse CD14 rat IgG2a and after washing with PBS, with anti-rat IgG-Alexa Fluor 647 (30 min, 4 °C each). The cells were then fixed, incubated with 4 μg/ml Hoechst 33342 as above, and subjected to confocal microscopy.

The cells were examined under an inverted Axio Observer Z.1 microscope (Zeiss) with a CSU-X1 spinning-disc unit (Yokogawa) equipped with a 63x oil immersion objective (NA 1.4) and an Evolve 512 EMCCD camera (Photometrics). Fluorescence was excited using 405- and 647-nm diode lasers and detected in the spectral range of 419-465 nm and 668-726 nm. Laser power and camera gain were adjusted to cover a 16-bit dynamic range. Optical sections were acquired using image format of 512 x 512 pixels and a voxel size of 0.212 μm in the x-y and 0.3 μm in the z-dimension. To determine the number of vesicles containing CD14 per cell, the obtained z-stacks were analyzed using Cell-Profiler 4.2. software (Broad Institute) [43]. To this end, images were converted to maximum z-projections and exported to a tiff format in ZEN 3.3 (Zeiss) software and processed in CellProfiler as follows. A median filter was applied, then nuclei were segmented basing on Hoechst 33342 fluorescence using the Otsu algorithm, and CD14-containing vesicles basing on Alexa Fluor 647 fluorescence using a manually adjusted threshold. The area covered by a particular cell was determined by expanding the nucleus mask by 40 pixels in each direction. The number of CD14-containing vesicles in a particular cell was thereby determined. To determine the amount of CD14 on the cell surface, z-stacks were converted to sum z-projections, and the average raw integral density per cell was calculated using ImageJ program [44] and a manually adjusted threshold.

### CD14 localization with immunofluorescence microscopy

Raw264 cells were seeded onto coverslips (3×10^4^ per 15×15 mm coverslip), grown for 24 h, washed in PBS and fixed in 3.7% formaldehyde in APHEM (60 mM Pipes, 25 mM HEPES, 10 mM EGTA, 4 mM MgCl_2_, pH 6.9). Subsequently, the cells were incubated with 15 mM NH_4_Cl in APHEM (10 min, room temp.), permeabilized with 0.005% digitonin in APHEM (10 min, room temp.) and blocked with 6% BSA in Tris-buffered saline (TBS; 30 min, room temp.). Some samples were additionally incubated with 2.5 µg/ml of a fusion protein containing the pleckstrin homology (PH) domain of human FAPP1 and the GST tag (FAPP1-PH-GST), [45] in 0.2% BSA/TBS for 60 min at room temp. and washed (five times x 5 min in 0.2% BSA/TBS). The cells were then incubated overnight with rat anti-CD14 IgG (clone Sa14-2; BioLegend) and goat anti-GST IgG (Rockland) or rabbit anti-Rab11 IgG or rabbit anti-golgin-97 IgG or rabbit anti-EEA1 IgG (all rabbit IgG from Cell Signaling Technology) in 0.2% BSA/TBS and, after washing as above, were exposed to appropriate secondary IgG: chicken anti-rat IgG-Alexa Fluor 647 (Invitrogen), donkey anti-goat IgG-FITC (Rockland) or donkey anti-rabbit IgG-FITC (Jackson ImmunoResearch) and 2 µg/ml Hoechst 33342 in 0.2 % BSA/TBS for 60 min at room temp. After washing, samples were mounted in mowiol/DABCO and examined under an LSM800 inverted confocal microscope (Zeiss) using a 63x oil objective (NA 1.4) with scan frame 553×553, speed 9, zoom 2.6, line averaging 2, pixel size 70 nm, and z interval 230 nm. Triple-stained images were obtained by sequential scanning for each channel to eliminate the crosstalk of the chromophores and to ensure a reliable quantification of colocalization. FITC fluorescence was excited at 488 nm, Alexa Fluor 647 at 640 nm, and Hoechst at 405 nm, and detected at 485-540, 637-700, and 400-487 nm, respectively. 16-bit z-stacks were used to build a 3D structure in Imaris 9.1.2. (Oxford Instruments) software. Regions of interest (ROI) related to peri-nuclear structures defined by staining with FAPP1-PH, golgin-97 or Rab11, and to EEA1-positive vesicles were selected using the surface detection algorithm of Imaris with a manually adjusted threshold. The sum of voxel intensities of CD14 staining and of the marker proteins [46] and also Pearson’s correlation coefficient (PCC) for quantifying the protein colocalization were calculated in the ROI. For a complete colocalization, PCC has the maximal value of 1. As found earlier [46], the values of the sum of voxel intensities did not follow a Gaussian distribution, therefore the median was used as a measure of average intensities and the nonparametric Mann-Whitney test to evaluate statistical significance. To measure the total intensity of CD14 staining in the cell, the z-stacks were converted to sum z-projections and raw integral intensity per cell was calculated using the ImageJ Analyze Particle feature and a manually adjusted threshold.

### Flow cytometry

Raw264 cells were seeded at 1×10^6^/well in 12-well culture plates, grown overnight in DMEM/10% FBS, suspended in ice-cold PBS and incubated in 2% mouse serum (Jackson ImmunoReserach) containing 0.01% NaN_3_ (30 min, 4°C). The cell-surface CD14 was labeled using rat anti-mouse CD14 IgG2a conjugated with FITC (clone Sa2-8; eBioscience) for 30 min at 4 °C (in the darkness). After that, cells were washed twice with ice-cold PBS and fixed with 3% formaldehyde. In control samples, the anti-CD14 IgG2a was omitted. After washing with PBS, cells were resuspended in PBS and their fluorescence was determined with a Becton Dickinson FACS Calibur flow cytometer. FITC fluorescence was detected using a 530/30 nm filter and an FL-1 detector. Cell debris was gated out by establishing a region around the population of interest on a Forward Scatter (FSC) versus Side Scatter (SSC) dot plot. Data were analyzed using BD CellQuest Pro software (BD Biosciences) and the amount of the cell-surface CD14 was calculated based on the geometric mean of fluorescence intensity.

### RNA isolation and RT-qPCR

RNA was isolated from cells using a Universal RNA purification kit (EUR_x_) and reverse-transcribed into cDNA using the High Capacity cDNA Reverse Transcription Kit (Thermo Fisher Scientific) according to the manufacturers’ instructions. The qPCR analysis was performed in a StepOnePlus instrument using Fast SYBR Green Master Mix (Thermo Fisher Scientific). Primers used are shown in Table 2. PCR conditions were the following: initial denaturation for 20 s at 95 °C followed by 40 cycles comprised of denaturation for 3 s at 95 °C and annealing/extension for 30 s at 60 °C. mRNA expression levels for investigated genes (relative to the mRNA level for the *Tbp* or *Hprt* gene, each variant run in triplicate) were calculated by the ΔΔC_T_ method.

### Cell fractionation

Cells (1.5×10^6^ per sample) were grown overnight and subjected to fractionation essentially as described earlier [25]. Briefly, cells were lysed in 500 μl of 0.1% Triton X-100, centrifuged (10 000 x *g*, 5 min, 4 °C), supernatants collected as Triton X-100-soluble fraction while pellets were solubilized in 500 μl of 1.8% octyl β-D-glucoside and centrifuged as above. Supernatants were collected as the DRM fraction, while the remaining pellets were solubilized in 500 μl of 4% SDS. Equal volumes of the fractions were subjected to 10% SDS-PAGE.

### Immunoblotting

To obtain whole cell lysates, cells (1.2×10^5^/well in 48-well plate) were rinsed with ice-cold PBS and lysed in 40 μl of a buffer containing 0.5% Triton X-100, 100 mM NaCl, 30 mM Hepes, 2 mM EDTA, 2 mM EGTA, protease and phosphatase inhibitors described above, pH 7.4. Protein concentration in the lysates was measured with BradfordUltra (Expedeon), the lysates were supplemented with sample buffer and subjected to 10% SDS-PAGE. After transfer onto nitrocellulose, the membrane was blocked and incubated with antibodies indicated in Table 3. Immunoreactive bands were visualized with chemiluminescence using SuperSignal WestPico substrate (Thermo Fisher Scientific) and analyzed densitometrically using the ImageJ program. Samples from the biotinylation assay and cell fractioning were blotted with appropriate primary antibodies followed by secondary antibodies conjugated with near-infrared fluorescent dyes IRDye 800CW or 680RD shown in Table 3 (LI-COR), and analyzed using an Odyssey M scanner (LI-COR) and Empiria Studio 2.2 software.

**Table 3.** List of used antibodies.

| Specificity of antibody | Host | Company | Catalog No. | Dilution | Application |
| --- | --- | --- | --- | --- | --- |
| Primary antibodies |  |  |  |  |  |
| flotillin-1 monoclonal IgG | Rabbit | Cell Signalling Techn. | #18634 | 1:1000 | IB |
| flotillin-2 monoclonal IgG | Rabbit | Cell Signalling Techn. | #3436 | 1:1000 | IB |
| Jak1 monoclonal IgG | Rabbit | Cell Signalling Techn. | #3344 | 1:1000 | IB |
| TNFα monoclonal IgG | Rabbit | Cell Signalling Techn. | #11948 | 1:1000 | IB |
| CD14 monoclonal IgG | Rat | BD Pharmingen | #553738 | 1:1000 | IB |
| TLR4 monoclonal IgG | Rabbit | Cell Signalling Techn. | #14358 | 1:1000 | IB |
| actin monoclonal IgG | Mouse | BD Biosciences | # 612656 | 1:30 000 | IB |
| phospho-IκBα monoclonal IgG | Rabbit | Cell Signalling Techn. | #2859 | 1:1000 | IB |
| phospho-IRF3 monoclonal IgG | Rabbit | Cell Signalling Techn. | #4947 | 1:1000 | IB |
| CD14 monoclonal IgG2a-FITC | Rat | eBioscience | #11-0141-82 | 1:200 | FC |
| CD14 monoclonal IgG2a | Rat | BioLegend | #123302 | 1:300 | RA, IF |
| IgG2a | Rat | BioLegend | #400502 | 1:300 | RA |
| GST IgG | Goat | Rockland | #200-101-200 | 1:300 | IF |
| EEA1 IgG | Rabbit | Cell Signalling Techn. | #3288 | 1:200 | IF |
| Rab11 IgG | Rabbit | Cell Signalling Techn. | #5589 | 1:200 | IF |
| golgin-97 IgG | Rabbit | Cell Signalling Techn. | #13192 | 1:200 | IF |
| Secondary antibodies |  |  |  |  |  |
| mouse IgG-HRP | Goat | Jackson ImmunoResearch | # 115-035-003 | 1:30 000 | IB |
| rabbit IgG-HRP | Goat | Rockland | # 611-1302 | 1:5000-1:10 000 | IB |
| rat IgG-HRP | Goat | Merck | # A9037 | 1:8000 | IB |
| rat IgG-IRDye 800CW | Goat | LI-COR | #926-32219 | 1:10000 | IB-LICOR |
| mouse IgG-IRDye 800CW | Goat | LI-COR | #952-32210 | 1:10000 | IB-LICOR |
| rabbit IgG-IRDye 680RD | Goat | LI-COR | #925-68071 | 1:10000 | IB-LICOR |
| rat IgG-Alexa Fluor 647 | Chicken | Invitrogen | #A-21472 | 1:300-1:500 | RA, IF |
| rabbit IgG-FITC | Donkey | Jackson ImmunoResearch | #711-095-152 | 1:300 | IF |
| goat IgG-FITC | Donkey | Rockland | #605-702-125 | 1:300 | IF |

IB, immunoblotting; FC, flow cytometry; IF, immunofluorescence; RA, recycling assay

### Data analysis

The significance of differences was calculated using Student’s *t-*test or 2-way ANOVA and Scheffe’s or Tukey’s post hoc test; for non-normal variables Mann-Whitney test or Kruskal-Wallis test and Dunn’s post hoc were used. The data analysis was performed using the JASP software environment. The statistical tests used in experiments are given in Figure legends. *P*-values < 0.05 were defined as statistically significant and are shown in Figures. Some data are presented as boxplots, where the box encompasses the first and third quartiles with a line through the median. The top and bottom whiskers denote the maximum and minimum data points, respectively, excluding data 1.5 times the interquartile range (IQR) which are denoted as points.

## Results

### Silencing of *Flot-2* with shRNA leads to depletion of flotillin-2 and flotillin-1 in Raw264 cells

To assess the role of flotillins in LPS-induced signaling we aimed at obtaining macrophage cells with a stably silenced expression of respective gene(s). An RT-qPCR analysis indicated that the amount of flotillin-2 mRNA in Raw264 macrophage-like cells exceeded that of flotillin-1 about 4.2-fold. Also in J774 macrophage-like cells and in macrophages isolated from mouse peritoneum flotillin-2 mRNA prevailed over that of flotillin-1, 1.8- and 3.5-fold, respectively (Fig. 1A, B). Raw264 and J774 cells were relatively rich in CD14 mRNA in comparison to primary macrophages, while th TLR4 mRNA level was the highest in the latter cells (Fig. 1C, D). Since the expression of *Flot2* dominated over *Flot1* in all the cells tested, we silenced the expression of *Flot2* in Raw264 cells and used them for further studies.

**Fig. 1.**
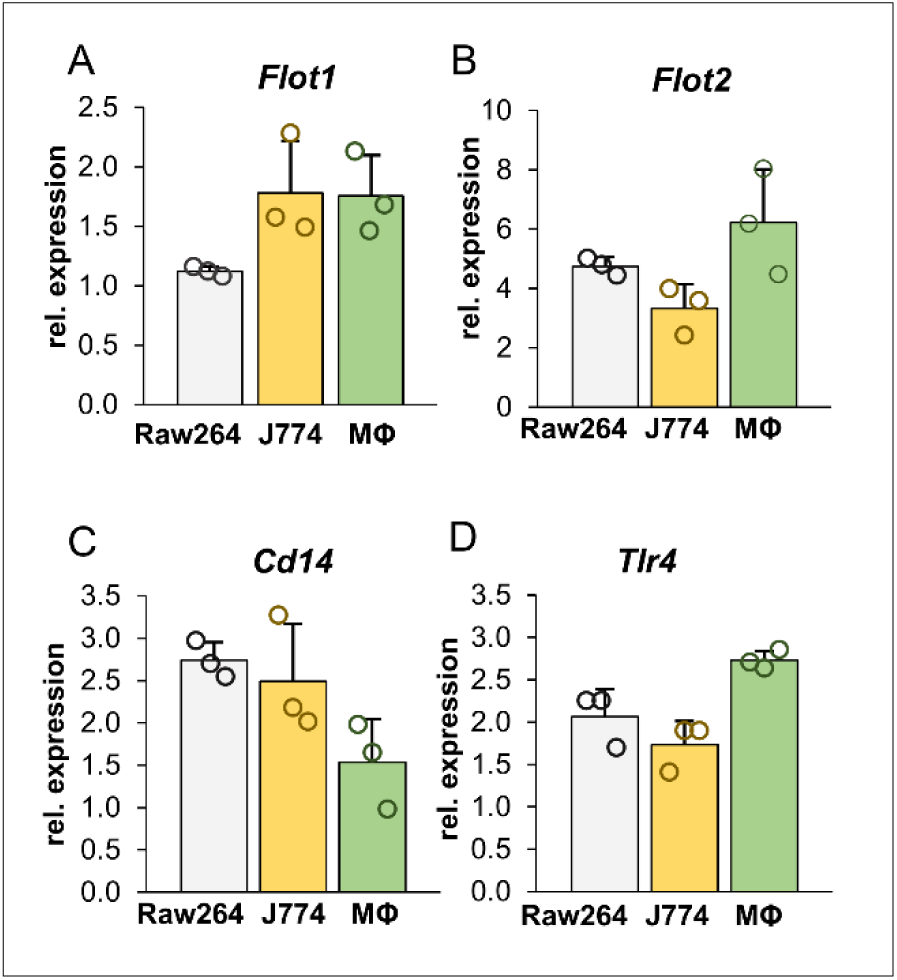
mRNA levels of flotillin-1, flotillin-2, CD14, and TLR4 in Raw264 and J774 cells, and mouse peritoneal macrophages (MΦ). (A, B, D) *Flot1* (A)*, Flot2* (B), and *Tlr4* (D) transcripts were quantified by RT-qPCR relative to *Tbp* while that of *Cd14* (C) relative to *Hprt* due to the comparable abundance of those transcripts. Data shown are mean ± SD from three experiments.

For this purpose, Raw264 cells were transfected separately with five different lentiviral particles bearing five different shRNA sequences and twice with control shRNA bearing a non-mammalian shRNA. An RT-qPCR analysis revealed that four of the shRNAs used effectively silenced *Flot2*, expression with the shRNA variant No. 2 being the exception (Supplementary Fig. 1A). On average, the four effective shRNA species gave a reduction of the relative *Flot2* mRNA level by 76% compared to the control cells transfected with the non-mammalian shRNAs, with most profound reduction, reaching 81%, observed for shRNA No. 5 (Fig. 2A, Supplementary Fig. 1A). Immunoblotting revealed that small amounts of flotillin-2 protein (on average about 12%, with only 4% in variant No. 5) were still present in the four Raw264 transformants (Fig. 2B-D, Supplementary Fig.1B). Further experiments were performed using transformants No. 1, 3 and 5, all obtained with shRNA species targeting the 3’-untranslated region of flotillin-2 mRNA (Table 1). We found, that these transformed cells were also depleted of flotillin-1 protein, and the extent of that depletion correlated with the reduction of flotillin-2 protein (Fig. 2B-D), in agreement with earlier studies [47, 48]. Also, the flotillin-1 mRNA level was reduced significantly, by about 7, 15 and 34% in the transformant lines 1, 3 and 5, respectively (Fig. 2E). Because we aimed to examine the effect of flotillin depletion on the response of the cells to LPS, we additionally determined the levels of CD14 and TLR4 mRNAs in those cells; they were decreased as well, on average by about 21% and 15%, respectively (Fig. 2F, G).

**Fig. 2.**
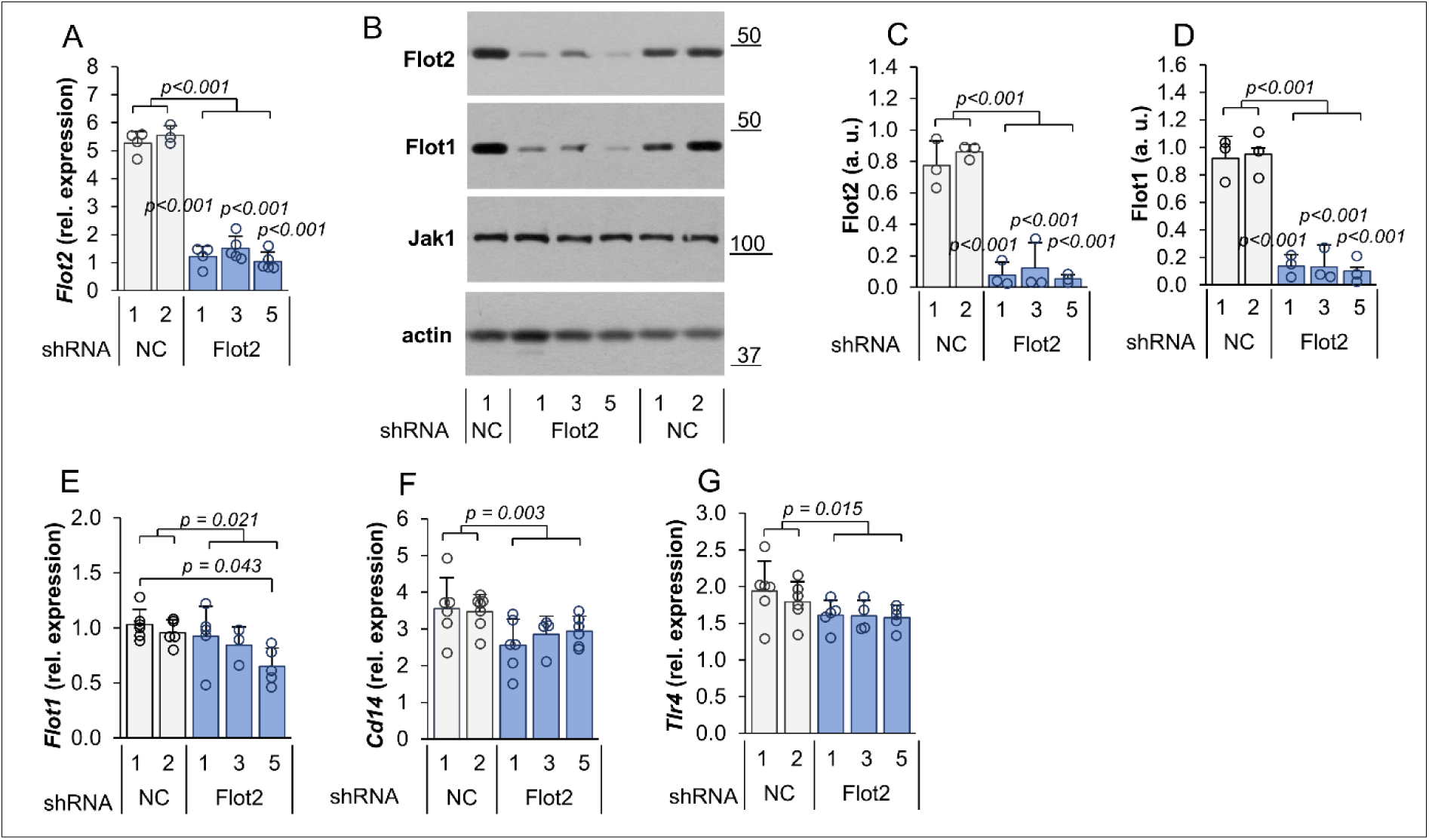
The effect of *Flot2* knock-down on the expression of selected genes and flotillin-1 and - 2 levels in Raw264 cells. Cells were transfected with *Flot2*-specific shRNA variants No. 1, 3, 5 (Flot2 shRNA) or control shRNA (NC1, NC2 shRNA). (A, E-G) RT-qPCR analysis of *Flot2* (A), *Flot1* (E), *Cd14* (F) and *Tlr4* (G) transcripts quantified as in Fig. 1. (B-D) The abundance of flotillin-1 and -2 determined by immunoblotting (B) followed by densitometric analysis (C, D). The flotillins were quantified in indicated cells relative to actin. Molecular weight markers are shown on the right in kDa. Jak1 and actin were visualized to verify equal loading of protein between wells. Data shown are mean ± SD from at least three experiments. Significantly different values as indicated by 1-way ANOVA with Scheffe’s post hoc test are marked. In (A, C, D) the *p* values for individual Flot2 shRNA variants were < 0.001 relative to both NC variants

### Depletion of flotillins reduces the CD14 level and inhibits LPS-induced signaling in Raw264 cells

To determine whether the flotillin depletion affected LPS-induced signaling, cells were stimulated with 10 or 100 ng/ml LPS for 1 h and the level of selected proteins of interest was examined by immunoblotting. The depletion of flotillins seen in unstimulated cells did not change upon their stimulation (Fig. 3A-C). The level of CD14 protein was reduced substantially, by about 35%, in resting cells. Upon stimulation with LPS, the amount of CD14 protein rose by about 14 and 25% (10 or 100 ng/ml LPS) in control cells and moderately also in the transformants. Ultimately, however, the level of CD14 in LPS-stimulated cells remained lower in the flotillin-depleted cells than in controls, by 28% and 42%, respectively (Fig. 3A, D).

**Fig. 3.**
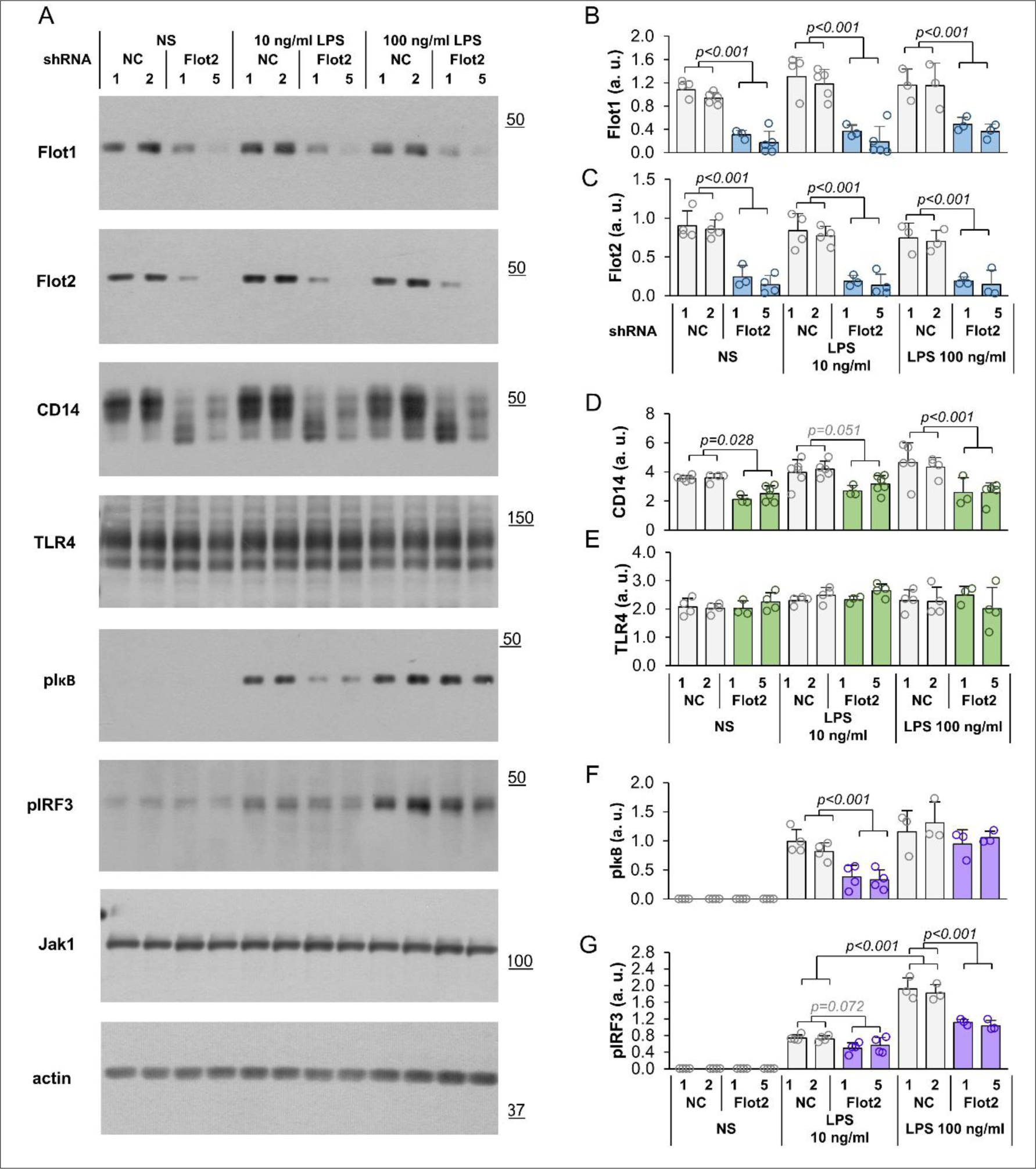
The effect of *Flot2* knock-down on the level of proteins involved in LPS-induced signaling. Cells were transfected with *Flot2*-specific shRNA variants No. 1 or 5 (shRNA Flot2) or control shRNA (NC1, NC2 shRNA). Cells were left unstimulated (NS) or were stimulated with 10 ng/ml or 100 ng/ml LPS (1 h, 37 °C). (A) Immunoblotting analysis of the abundance of indicated proteins in the cells. Molecular weight markers are shown on the right in kDa. Jak1 and actin were visualized to verify equal loading of protein between wells. (B-E) Densitometric analysis of (B) flotillin-1, (C) flotillin-2, (D) CD14, (E) TLR4, (F) phosphorylated IκB (pIκB), (G) phosphorylated IRF3 (pIRF3) content relative to Jak1. Data shown are mean ± SD from at least three experiments. Significantly different values as indicated by 1-way ANOVA with Scheffe’s post hoc test are marked

In striking contrast to CD14, the level of TLR4 in flotillin-depleted resting cells did not differ significantly from the control level, and the 1-h stimulation with LPS did not affect it in any of those cells either (Fig. 3A, E). Despite that, the deficiency of flotillins reduced the TLR4-dependent activation of two major transcription factors, NFκB and IRF3, triggered by LPS via, respectively, both the MyD88- and TRIF-dependent routes and the TRIF-dependent one exclusively [9, 14]. Notably, the activation of NFκB, indicated by phosphorylation of its regulatory subunit IκB, was reduced significantly only in flotillin-depleted cells stimulated with 10 ng/ml LPS, on average by 64% *vs.* control cells, while at 100 ng/ml LPS the difference between flotillin-depleted and control cells (14% reduction) was non-significant. The IκB phosphorylation in controls was only slightly higher at 100 than at 10 ng/ml LPS (Fig. 3A, F). In contrast, the intensity of the TRIF-dependent phosphorylation of IRF3 in LPS-stimulated control cells correlated significantly with the LPS concentration used, as did the extant of its reduction in the flotillin-depleted cells, which amounted to 28% at 10 ng/ml and as much as 43% at 100 ng/ml LPS (Fig. 3A, G).

Taken together, the data show a negative influence of flotillin depletion on the abundance of CD14 but not of TLR4 in cells, and a resulting clear-cut reduction of the TRIF-dependent signaling of TLR4 with the MyD88-dependent one affected significantly at the lower LPS concentration only.

### Depletion of flotillins limits LPS-induced production of cytokines

To determine the ultimate impact of the flotillin depletion on the pro-inflammatory responses of Raw264 cells, we determined the amounts of TNFα and CCL5/RANTES, two cytokines produced, respectively, mainly in the MyD88-dependent and strictly in the TRIF-dependent manner [11]. In control cells, the production of TNFα was about 2-fold higher at 10 than at 100 ng/ml LPS (Fig. 4A), and flotillin depletion affected it only in cells stimulated with 10 ng/ml LPS; it was lower in transformants No. 1, 2 and 3 by 31-46% of the control level. In contrast, the CCL5/RANTES production was 1.5-fold higher in control cells stimulated with 100 ng/ml LPS than at 10 ng/ml LPS. In the flotillin-depleted cells it was lower by 36-58% upon stimulation with 10 ng/ml LPS and by at least 21% (37% in transformants No. 5) at 100 ng/ml LPS (Fig. 4B). Thus, the present results obtained using ELISA, were consistent with the results of immunoblotting analysis of IκB and IRF3 phosphorylation shown in Fig. 3.

**Fig. 4.**
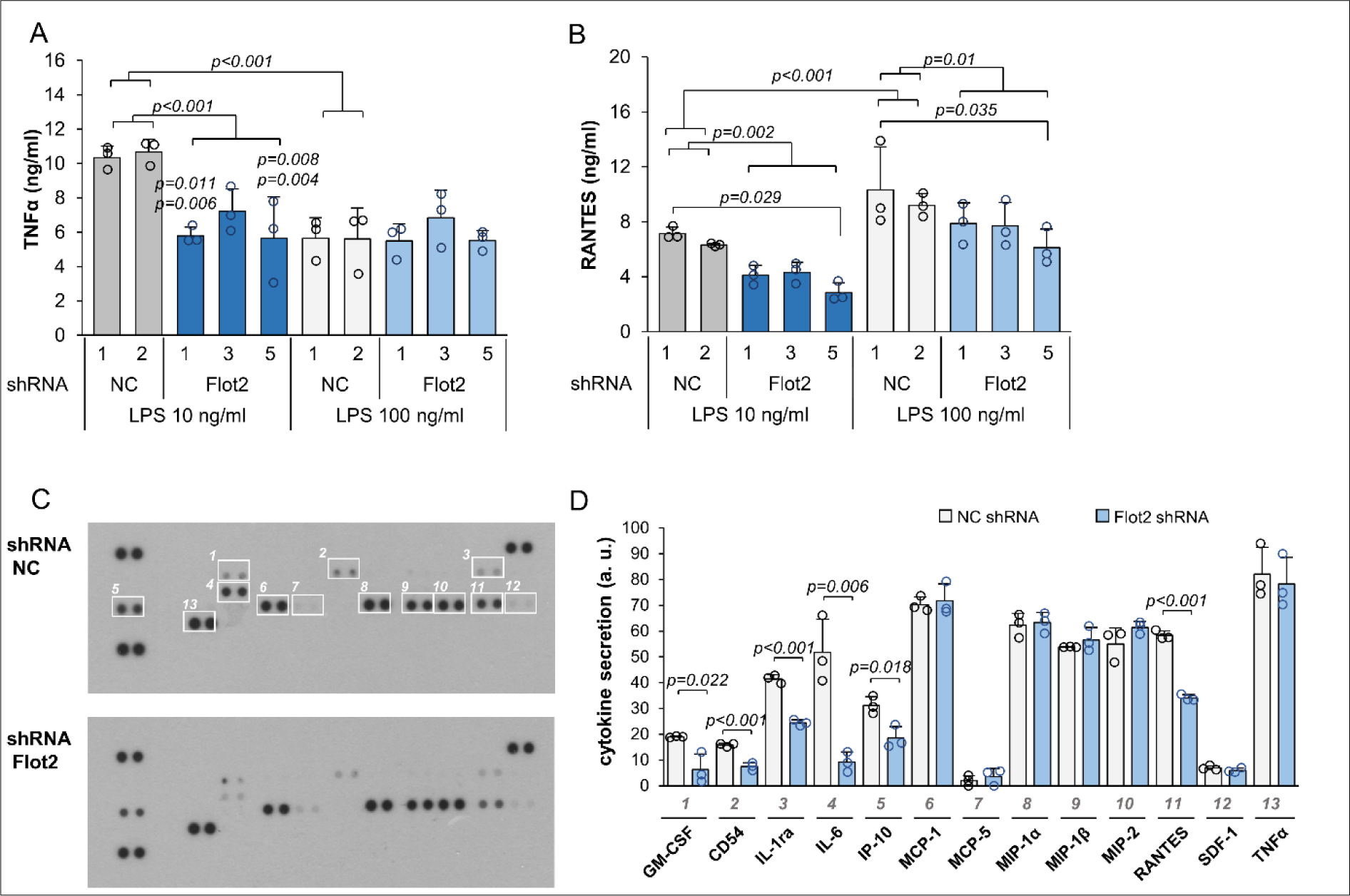
Inhibition of LPS-induced cytokine production by the flotillin depletion. Cells were transfected with *Flot2*-specific shRNA variants No. 1, 3, 5 (Flot2 shRNA) or control shRNA (NC1, NC2 shRNA). (A, B) The concentration of TNFα (A) and CCL5/RANTES (B) was determined in culture supernatants with ELISA. Cells were left unstimulated or were stimulated with 10 ng/ml or 100 ng/ml LPS (6 h, 37 °C). No cytokine production was found without cell stimulation. Data shown are mean ± SD from three experiments. Significantly different values as indicated by 1-way ANOVA with Tukey’s post hoc test are marked. (C, D) Cytokine array detection (C) followed by densitometric analysis of immunoblots (D). Dots left unmarked in the upper blot in (C) served as internal standards used to normalize signals between membranes. Cells were stimulated with 10 ng/ml LPS (4 h, 37 °C). Results for cells transfected with *Flot2*-specific shRNA variant No. 5 and control shRNA NC1 are shown. Data shown in (D) are mean ± SD from three experiments. Significantly different values as indicated by Student’s *t*-test are marked

To get a more complete picture of how the LPS-induced cytokine production is affected by flotillin deficiency, we analyzed the production of an array of inflammatory markers by cells stimulated with 10 ng/ml LPS (Fig. 4C, D). Altogether, 13 inflammatory markers were secreted by Raw264 cells in these conditions. Among them, the secretion of IP-10, IL-1ra, CCL5/RANTES, GM-CSF, IL-6, and CD54/ICAM-1 was significantly lower for the flotillin-depleted cells (Fig. 4C, D). Notably, the production of IP-10, IL-1ra, and CCL5/RANTES depends strictly on the involvement of TRIF and IRF3 activation, and that of IL-6 and CD54/ICAM-1 has also been reported to depend strongly on this TLR4-triggered signaling pathway [14, 15, 49–52]. The production of those inflammatory markers was reduced by 40-66% and by as much as 82% in the case of IL-6 (Fig 4C, D). Most of those factors have a pro-inflammatory activity except for IL-1ra, which is anti-inflammatory cytokine and can negatively regulate the pathogenesis of inflammation [52]. On the other hand, the assay revealed no effect of flotillin deficiency on the level of cytokines produced in the MyD88-dependent pathway of TLR4, including the MIP family chemokines and TNFα (Fig. 4C, D), the latter in contrast to the ELISA results presented above (see Fig. 4A). This discrepancy could result from differences in the sensitivity and the mode of cytokine detection between these two assays. The cytokine array seems less suitable for assessing the level of relatively abundant cytokines, such as the MyD88-dependent ones, due to the high, saturated optical density of respective spots.

To sum up, the obtained data indicate that the participation of flotillins is required for a maximal LPS-induced production of cytokines. The knock-down of flotillins affected particularly the TRIF-dependent signaling of TLR4, and an effect on the MyD88-dependent pathway also seems possible, especially in cells stimulated with a low LPS concentration (10 ng/ml). Thus, the involvement of flotillins in LPS-induced signaling is manifested in conditions that require the engagement of CD14 for TLR4 activation and endocytosis [16, 18, 53]. For this reason, we decided to establish whether flotillins take part in determining the cell surface level of CD14 known to condition the extent of the TRIF-dependent signaling of TLR4 [22].

### Flotillin depletion reduces the surface level of CD14

By using flow cytometry to determine the abundance of cell-surface CD14 we found that flotillin depletion in Raw264 cells decreased the surface pool of CD14 by about 26% in comparison with control cells (Fig. 5A). Using cell fractionation into Triton X-100-soluble and -insoluble (DRM) fractions, we found that flotillin depletion strongly reduced the amount of the mature form of CD14 (CD14 bands with slower gel migration) found in the DRM fraction despite a relatively modest content of flotillin-2 itself in this fraction in control cells (Fig. 5B). In contrast, another raft-associated protein, Lyn kinase, was present in comparable amounts in the DRM fraction in control and flotillin-depleted cells (Fig. 5B), indicating the specificity of the effect of flotillin depletion on raft-associated CD14. As expected, transferrin receptor was found exclusively in the Triton X-100-soluble fraction regardless of the flotillin status of the cells (Fig. 5B). This fraction contained also the immature forms of CD14, substantial amounts of flotillin-2 and actin, while small amounts of the examined proteins were found in the SDS-soluble (cytoskeletal) fraction (Fig. 5B).

**Fig. 5.**
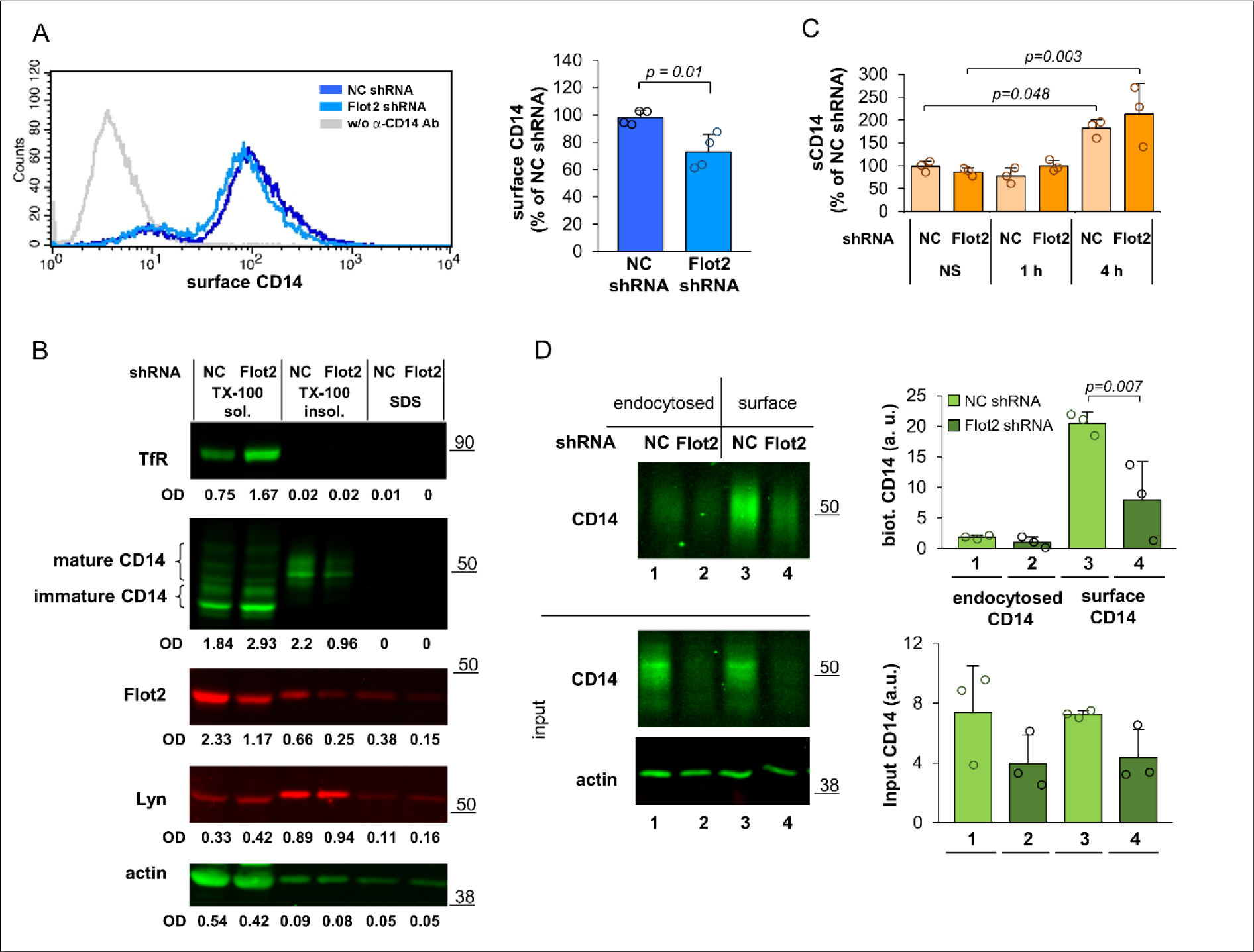
The effect of *Flot2* knock-down on CD14. Cells were transfected with *Flot2*-specific shRNA variant No. 5 (Flot2 shRNA) or control shRNA NC1 (NC shRNA). (A) The surface level of CD14 determined by flow cytometry (left) and expressed as percentage of the value in control cells (right). Data shown are mean ± SD from four experiments. (B) Fractionation of Tritonx-100 lysates of the cells. TX-100 sol. - fraction soluble in 0.1% Triton X-100, TX-100 insol. - fraction insoluble in Triton X-100, SDS - fraction soluble in 4% SDS. Numbers below blots show the relative content of the respective protein in each fraction determined by densitometry (OD). Results of one representative experiment of two run in duplicate are shown. (C) The content of sCD14 in culture supernatants determined with ELISA. Cells were left unstimulated (NS) or were stimulated with 10 ng/ml LPS for 1 or 4 h (37 °C). (D) Endocytosis of CD14 determined by a biotinylation-based assay. See text for details of the procedure. To avoid overloading, only 1/3 of the samples containing the cell surface CD14 was loaded on the gel. The total amount of CD14 relative to actin in input cell lysates is also shown (lower panels). Molecular weight markers are shown on the right in kDa. Data shown in (C, D) are mean ± SD from three experiments. Significantly different values as indicated by Student’s *t*-test (A) and 1-way ANOVA with Tukey’s post hoc test (C, D) are marked

Since the reduction of the cell surface (mature) CD14 could be caused by its enhanced shedding, we determined the amount of soluble CD14 (sCD14) released by control and flotillin-depleted cells. Flotillin depletion did not reduce the amount of sCD14 released by resting cells while in LPS-stimulated cells it was slightly higher than in control cells (Fig. 5C). Given the reduction of *Cd14* expression and the total cellular CD14 abundance, the release of sCD14 by may in fact have been more efficient in flotillin-depleted cells to reach and even surpass the control level.

Also, changes of CD14 endocytosis could affect its cell surface level, as CD14 is known to undergo a slow constitutive uptake in resting cells [21, 22]. To investigate this possibility, we labeled the cell surface proteins of Raw264 cells with a membrane-impermeable biotin derivative and compared the amounts of labeled CD14 still associated with the plasma membrane with that found in the cell interior 10 minutes later; the stronger was the CD14 endocytosis, the higher its fraction would be expected in the latter pool. This assay confirmed the reduction of the amount of CD14 present on the surface of flotillin-depleted cells (Fig. 5D, compare bars 3 and 4) and also a strong reduction of the total amount of the protein in input samples (Fig. 5D, compare bars 1a and 3a with 2a and 4a, respectively). According to this assay, the cell surface level of CD14 was diminished in flotillin-depleted cells by as much as 61% which exceeded the extent of the CD14 depletion detected on the surface of intact cells by flow cytometry (Fig. 5D, compare with Fig. 5A). We believe that this quantitative difference reflects the differences between the two assays run on intact *vs.* lysed cells. Ten minutes after the labeling reagent was washed off, 9% of the labeled CD14 was internalized in control cells (Fig. 5D, compare bars 1 and 3), and about 13% in those depleted of flotillins Fig. 5D, compare bars 2 and 4). These data indicate that the depletion of flotillins in Raw264 cells does not affect the rate of CD14 internalization in a significant manner, albeit its slight enhancement can be observed when related to the initial amount of CD14 on the cell surface.

### Flotillin depletion enhances CD14 recycling

The data presented above prompted us to analyze the influence of flotillin depletion on the recycling of CD14. For this purpose, we applied a microscopic assay developed by us recently [22] which is based on the labeling of cell-surface CD14 with specific antibodies. Following successive incubations of the cells with the anti-CD14 antibody and a secondary fluorescent antibody, separated by cell warming and removal of the CD14-bound antibodies remaining on the cell surface, only the recycling pool of CD14 is eventually visible in vesicles inside the cells (Fig. 6A). In the flotillin-depleted cells the number of vesicles containing recycling CD14 was significantly higher (about 2-fold) than in control cells (Fig. 6B and Supplementary Fig. 2A, compare lanes 3 and 1).

**Fig. 6.**
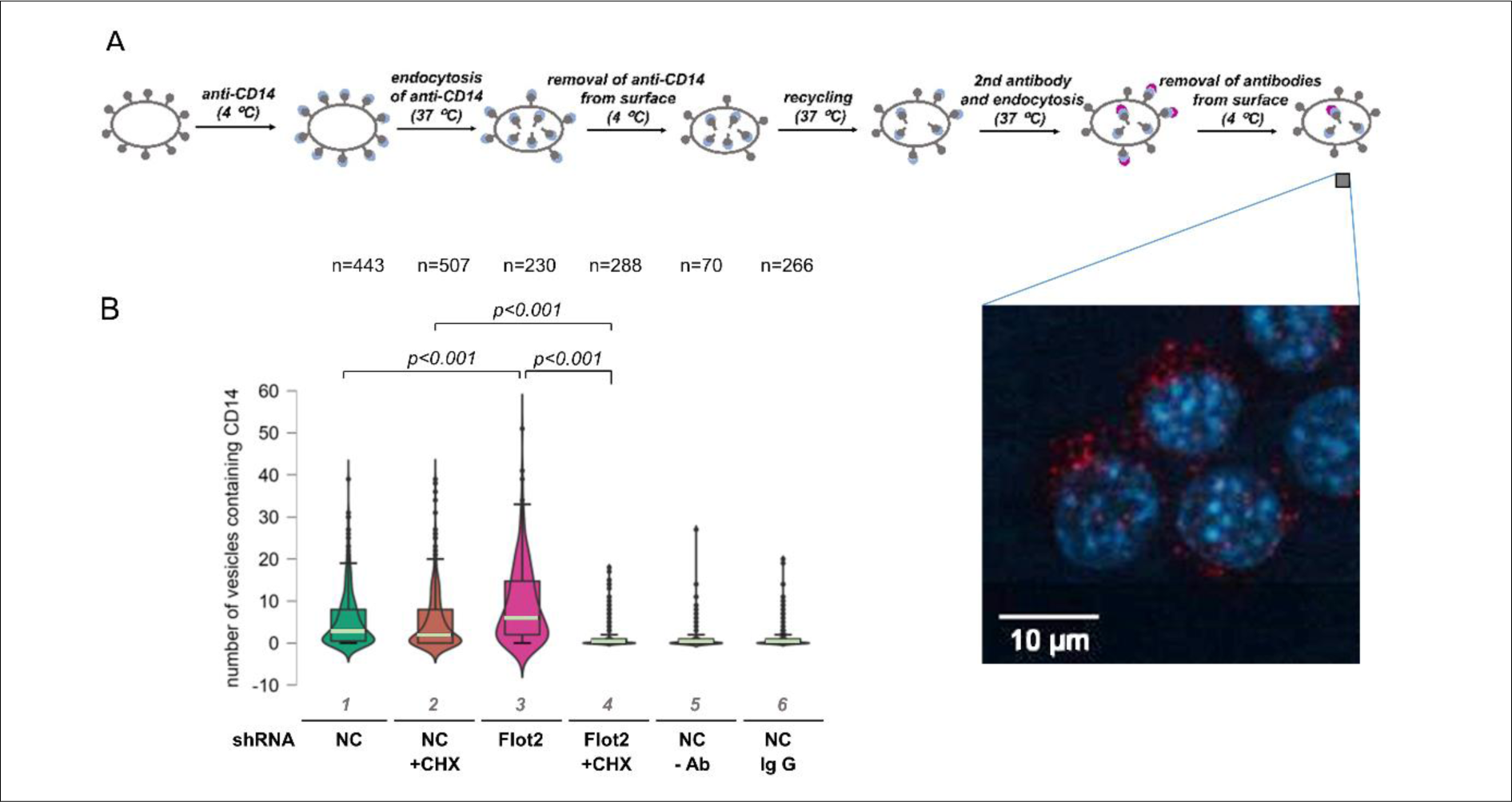
Enhancement of CD14 recycling in flotillin-depleted cells. Cells were transfected with *Flot2*-specific shRNA variant No. 5 (Flot2 shRNA) or control shRNA NC1 (NC shRNA). (A) Scheme of the fluorescent antibody-based assay of CD14 recycling with a micrograph showing CD14-containing vesicles (red) and nuclei (blue). (B) Quantitation of CD14 recycling. The cells were examined under a fluorescence microscope and vesicles containing fluorescently labeled CD14 representing the recycling CD14 were counted in *n* number of cells. When indicated, the cells were pretreated with 20 μg/mL CHX prior to the assay and CHX was also present during the assay. In control samples, rat isotype IgG2a was used instead of the CD14-specific antibody or the antibody was omitted. Box plots represent the median (light green lines) and 25th/75th quartiles of the number of vesicles containing fluorescently labeled CD14 per cell in one experiment of two. Significantly different values as indicated by Kruskal-Wallis test followed by Dunn’s multiple comparisons post-hoc test are marked Parenthetically, this assay further confirmed the reduction of the cell surface level of CD14 in flotillin-depleted cells (Supplementary Fig. 2B). To establish to what extent the recycling of CD14 in both cell types was affected by newly synthesized CD14, the cells were pretreated with 20 μg/ml cycloheximide (CHX) for 30 minutes prior to the assay. CHX decreased the rate of CD14 recycling in control cells (the median number of CD14-containing vesicles decreased by 1/3; Fig. 6B and Supplementary Fig. 2A, compare lanes 1 and 2) and unexpectedly abolished it fully in flotillin-depleted cells (Fig. 6B and Supplementary Fig. 2A, compare lanes 3 and 4). This indicated that in the flotillin-depleted cells only the newly synthesized protein CD14 underwent a single round of endocytosis and recycling following its initial transport to the plasma membrane. Moreover, taking into account the low surface level of CD14 in these cells and only an insignificantly increased rate of its endocytosis, the CD14 recycling *per se* seems to be strongly enhanced in the absence of flotillins.

### Flotillin depletion leads to accumulation of CD14 in EEA1- and golgin-97-positive compartments

We broadened the investigation of the CD14 recycling in flotillin-depleted cells by analyzing its distribution in cellular compartments involved in protein sorting/recycling, i.e., early endosomes marked by EEA1, Rab11-positive endocytic recycling compartments (ERC), and the *trans*-Golgi network (TGN) identified by the presence of golgin-97. For comparison, we also determined the distribution of CD14 in the Golgi apparatus which accommodates newly synthesized CD14 *en route* to the plasma membrane. As a marker for this compartment we used the PH domain of the FAPP1 protein which binds to the PI(4)P/Arf1 complex in the Golgi [45, 54, 55]. Optical sections of triple-labeled cells (including nucleus labeling) were used to reconstruct 3D images of studied structures which were identified on the basis of the presence of the marker proteins in combination with their cellular localization. Namely, the ERC, TGN, and Golgi were defined as perinuclear structures enriched in respective marker proteins, whereas early endosomes were seen as dispersed EEA1-positive vesicles. The amount of CD14 in each compartment was determined by calculating the sum of its voxel intensities in each cell via 3D image analysis [46] and related to the total fluorescent signal for CD14 in the sum z-projections of the same cell. The latter ratio reflected the fraction of CD14 localized in a given compartment. We also assessed the colocalization of CD14 with the respective protein markers of each compartment by calculating PCC, and additionally determined the sum of voxel intensities representing the marker protein to reveal whether flotillin depletion affected the compartment itself (Table 4).

**Table 4.**
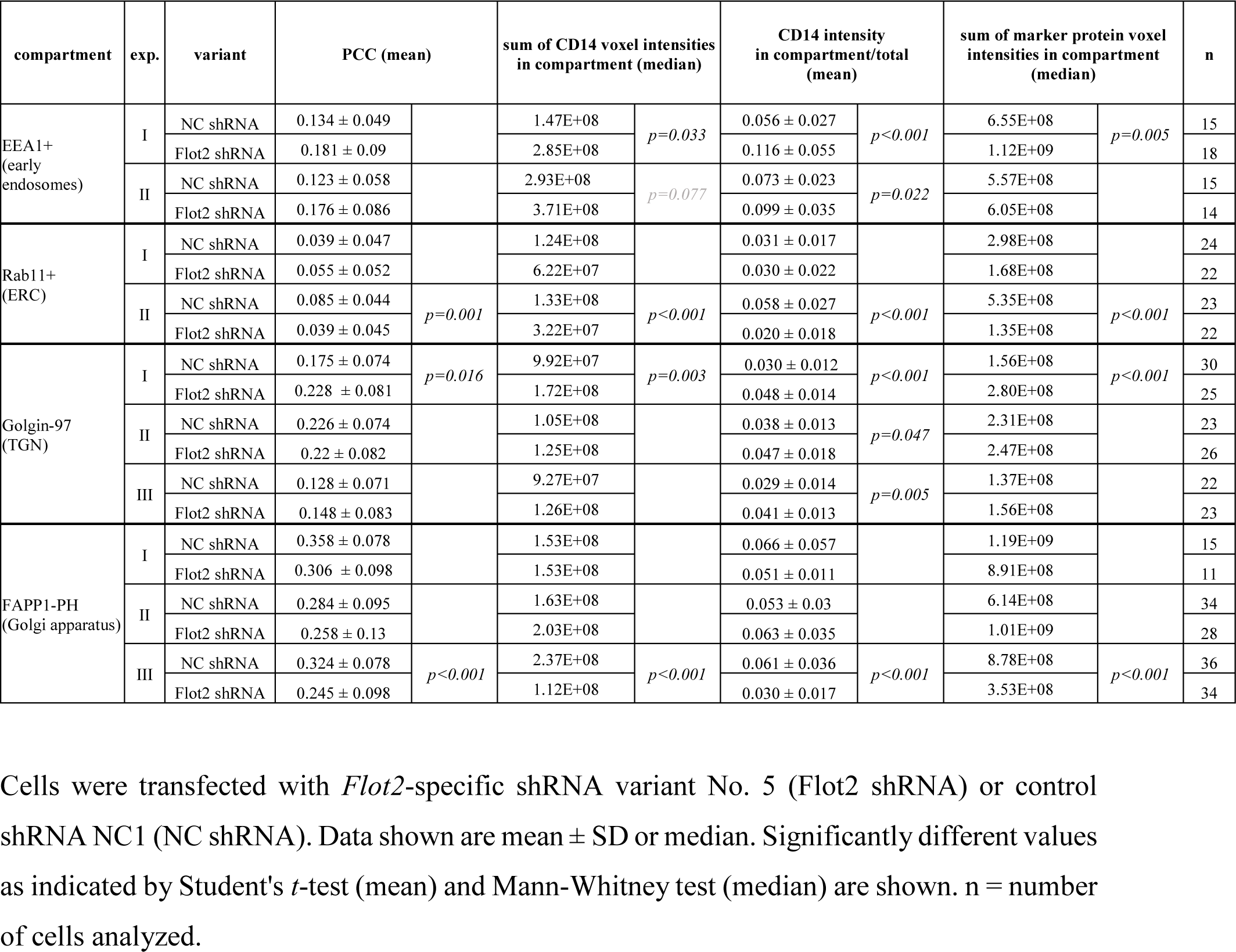
CD14 localization in cellular compartments involved in its trafficking.

| compartment | exp. | variant | PCC (mean) |  | sum of CD14 voxel intensities in compartment (median) |  | CD14 intensity in compartment/total (mean) |  | sum of marker protein voxel intensities in compartment (median) |  | n |
| --- | --- | --- | --- | --- | --- | --- | --- | --- | --- | --- | --- |
| EEA1+ (early endosomes) | I | NC shRNA | 0.134 ± 0.049 |  | 1.47E+08 | *p*=0.033 | 0.056 ± 0.027 | *p*<0.001 | 6.55E+08 | *p*=0.005 | 15 |
|  |  | Flot2 shRNA | 0.181 ± 0.09 |  | 2.85E+08 |  | 0.116 ± 0.055 |  | 1.12E+09 |  | 18 |
|  | II | NC shRNA | 0.123 ± 0.058 |  | 2.93E+08 | *p*=0.077 | 0.073 ± 0.023 | *p*=0.022 | 5.57E+08 |  | 15 |
|  |  | Flot2 shRNA | 0.176 ± 0.086 |  | 3.71E+08 |  | 0.099 ± 0.035 |  | 6.05E+08 |  | 14 |
| Rab11+ (ERC) | I | NC shRNA | 0.039 ± 0.047 |  | 1.24E+08 |  | 0.031 ± 0.017 |  | 2.98E+08 |  | 24 |
|  |  | Flot2 shRNA | 0.055 ± 0.052 |  | 6.22E+07 |  | 0.030 ± 0.022 |  | 1.68E+08 |  | 22 |
|  | II | NC shRNA | 0.085 ± 0.044 | *p*=0.001 | 1.33E+08 | *p*<0.001 | 0.058 ± 0.027 | *p*<0.001 | 5.35E+08 | *p*<0.001 | 23 |
|  |  | Flot2 shRNA | 0.039 ± 0.045 |  | 3.22E+07 |  | 0.020 ± 0.018 |  | 1.35E+08 |  | 22 |
| Golgin-97 (TGN) | I | NC shRNA | 0.175 ± 0.074 | *p*=0.016 | 9.92E+07 | *p*=0.003 | 0.030 ± 0.012 | *p*<0.001 | 1.56E+08 | *p*<0.001 | 30 |
|  |  | Flot2 shRNA | 0.228 ± 0.081 |  | 1.72E+08 |  | 0.048 ± 0.014 |  | 2.80E+08 |  | 25 |
|  | II | NC shRNA | 0.226 ± 0.074 |  | 1.05E+08 |  | 0.038 ± 0.013 | *p*=0.047 | 2.31E+08 |  | 23 |
|  |  | Flot2 shRNA | 0.22 ± 0.082 |  | 1.25E+08 |  | 0.047 ± 0.018 |  | 2.47E+08 |  | 26 |
|  | III | NC shRNA | 0.128 ± 0.071 |  | 9.27E+07 |  | 0.029 ± 0.014 | *p*=0.005 | 1.37E+08 |  | 22 |
|  |  | Flot2 shRNA | 0.148 ± 0.083 |  | 1.26E+08 |  | 0.041 ± 0.013 |  | 1.56E+08 |  | 23 |
| FAPP1-PH (Golgi apparatus) | I | NC shRNA | 0.358 ± 0.078 |  | 1.53E+08 |  | 0.066 ± 0.057 |  | 1.19E+09 |  | 15 |
|  |  | Flot2 shRNA | 0.306 ± 0.098 |  | 1.53E+08 |  | 0.051 ± 0.011 |  | 8.91E+08 |  | 11 |
|  | II | NC shRNA | 0.284 ± 0.095 |  | 1.63E+08 |  | 0.053 ± 0.03 |  | 6.14E+08 |  | 34 |
|  |  | Flot2 shRNA | 0.258 ± 0.13 |  | 2.03E+08 |  | 0.063 ± 0.035 |  | 1.01E+09 |  | 28 |
|  | III | NC shRNA | 0.324 ± 0.078 | *p*<0.001 | 2.37E+08 | *p*<0.001 | 0.061 ± 0.036 | *p*<0.001 | 8.78E+08 | *p*<0.001 | 36 |
|  |  | Flot2 shRNA | 0.245 ± 0.098 |  | 1.12E+08 |  | 0.030 ± 0.017 |  | 3.53E+08 |  | 34 |

Cells were transfected with *Flot2*-specific shRNA variant No. 5 (Flot2 shRNA) or control shRNA NC1 (NC shRNA). Data shown are mean ± SD or median. Significantly different values as indicated by Student's *t*-test (mean) and Mann-Whitney test (median) are shown. n = number of cells analyzed.

We found that the flotillin deficiency facilitated the accumulation of CD14 in EEA1-positive endosomes, which was reflected both by an increase of the median sum of CD14 voxel intensities and by an increased fraction (by 35% and 94% in two experiments) of CD14 found in this compartment (Fig. 7A-C, Table 4). The colocalization of CD14 and EEA1 was low (Table 4), as CD14 was visible inside vesicles decorated on their surface with EEA1 (Fig. 7A_4_, B_4_). It is worth noting that the median sum of EEA1 voxel intensities also tended to be increased in the flotillin-depleted cells (Table 4), suggesting that the early endosomal compartment was larger in those cells and accommodated more CD14 than in control cells.

**Fig. 7.**
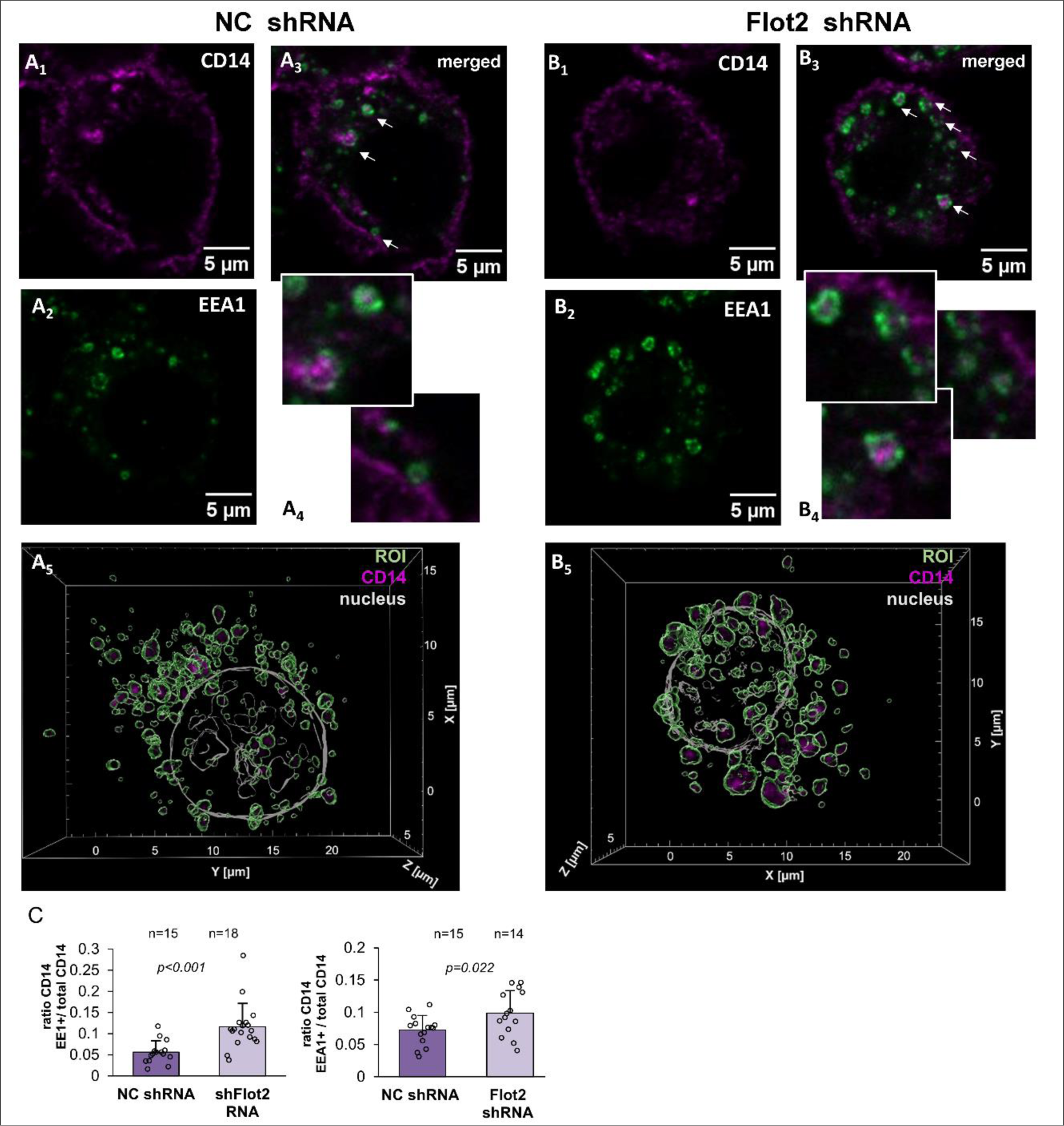
Enrichment of CD14 in EEA1-positive early endosomes in flotillin-depleted cells. (A) Raw264 cells transfected with control shRNA, variant NC1. (B) Cells transfected with shRNA specific against *Flot2,* variant No. 5. (A1, B1) Localization of CD14, (A2, B2) EEA1 and (A3, B3) merged images of CD14 and EEA1 localization. z-Stack images of three optical sections taken in the middle of a cell are shown. Colocalized CD14 and EEA1 appear as white spots. (A4, B4) Enlarged images of EEA1-positive endosomes indicated by arrows in (A3) and (B3) with CD14 visible in their interior. (A5, B5) Reconstructed 3D images of the EEA1-positive compartment delineated in green with CD14 visualized in pink. Contours of the nucleus detected by Hoechst 33342 staining are shown in grey. ROI as these were used for quantitative analysis of CD14 localization in the compartment - see Table 4. (C) Ratio of the fluorescence intensity of CD14 in EEA1-positive endosomes calculated as the sum of CD14 voxel intensities to the total CD14 fluorescence detected in z-projections of the respective cell. n, number of cells analyzed. Results of two independent experiments are shown. Significantly different values as indicated by Student’s *t*-test are marked

The enrichment of CD14 in early endosomes in the flotillin-depleted cells did not augment its localization in Rab11-positive ERC. On the contrary, all the ERC parameters analyzed were decreased or remained unaffected in those cells (Supplementary Fig. 3A-D, Table 4), suggesting that this compartment is not involved in the enhanced recycling of CD14. In marked contrast, the fraction of total CD14 found in the golgin-97-marked TGN was higher by up to 60% in the flotillin-depleted cells than in the control ones. In agreement, a moderate tendency for an increase of the median sum of CD14 voxel intensities in this compartment was observed in the flotillin-depleted cells (Fig. 8A-D, Table 4). Also the colocalization of CD14 with golgin-97 tended to be higher in those cells, albeit without a reproducible change in the median sum of golgin-97 voxel intensities itself (Fig. 8A-D, Table 4). These data indicate that the deficiency of flotillins facilitated CD14 recycling via the TGN rather than via the Rab11-positive compartment.

**Fig. 8.**
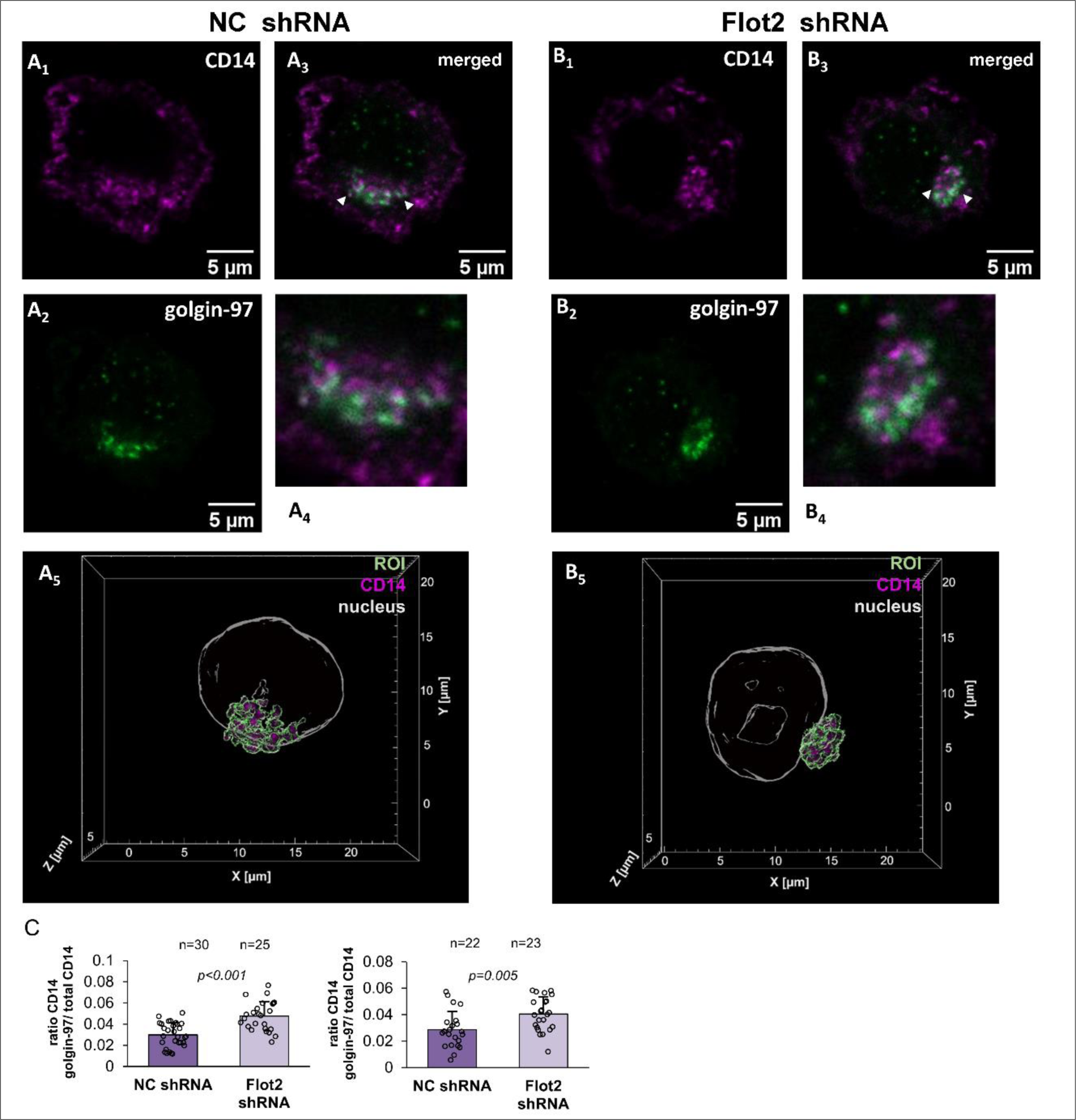
Enrichment of CD14 in golgin-97-positive TGN in flotillin-depleted cells. (A) Raw264 cells transfected with control shRNA, variant NC1. (B) Cells transfected with shRNA specific against *Flot2,* variant No. 5. (A1, B1) Localization of CD14, (A2, B2), golgin-97 and (A3, B3) merged images of CD14 and golgin-97 localization. z-Stack images of three optical sections taken in the middle of a cell are shown. Colocalized CD14 and EEA1 appear as white spots. (A4, B4) Enlarged images of golgin-97-positive TGN indicated by arrowheads in (A3) and (B3). (A5, B5) Reconstructed 3D images of the golgin-97-positive TGN delineated in green with CD14 visualized in pink. Contours of the nucleus detected by Hoechst 33342 staining are shown in grey. ROI as these were used for quantitative analysis of CD14 localization in the compartment - see Table 4. (C) Ratio of the fluorescence intensity of CD14 in golgin-97-positive TGN calculated as the sum of CD14 voxel intensities to the total CD14 fluorescence detected in z-projections of the respective cell. n, number of cells analyzed. Results of two independent experiments are shown. Significantly different values as indicated by Student’s *t*-test are marked

Finally, a fraction of CD14 was detected in the FAPP1-PH-decorated Golgi apparatus. Neither this fraction nor the median sum of CD14 voxel intensities in this compartment, nor the median sum of FAPP1-PH voxel intensities were consistently affected by the deficiency of flotillins (Supplementary Fig. 4A-D, Table 4). Interestingly, CD14 colocalized with FAPP1-PH to the highest degree among all the marker proteins investigated (PCC around 0.3), similarly in the flotillin-depleted and control cells (Table 4). These data indicate that flotillin depletion did not affect the anterograde transport of CD14 via the Golgi apparatus.

## Discussion

Flotillin-1 and -2 are ubiquitous scaffolding proteins associated with plasma membrane rafts and also with endo-membranes, affecting the signaling of plasma membrane receptors and their intracellular trafficking [27]. In this study we aimed to determine the role of flotillins in the CD14- and TLR4-mediated pro-inflammatory response of macrophages to LPS. For this purpose, we used shRNA transfection to obtain a set of Raw264 cells stably depleted of flotillin-2, finding that the cells were depleted of flotillin-1 as well. The phenomenon of a loss of the both flotillins after a directed down-regulation of one of them has already been reported and seems associated with the hetero-dimerization of flotillins increasing each other’s stability [32, 47, 56, 57]. Moreover, we also detected a down-regulation of flotillin-1 mRNA in Raw264 cells after *Flot2* silencing which could contribute to flotillin-1 depletion. Therefore, the reduction of LPS-induced inflammatory response of Raw264 cells described in this study needs to be ascribed to the depletion of the both flotillins even though some specific functions are attributed to flotillin-1 owing to its distinct *S-*palmitoylation pattern [56, 58].

We found that the depletion of flotillins diminished predominantly the TRIF-dependent endosomal signaling of TLR4 while the MyD88-dependent one was affected mainly in cells stimulated with the lower LPS concentration (10 ng/ml). One should note that the involvement of flotillins in LPS-induced signaling was manifested in conditions that required a CD14 involvement for TLR4 activation [7, 8, 16, 18, 53]. Indeed, a line of data obtained in this study indicates that flotillins modulate the LPS-induced signaling by affecting the abundance of CD14. The flotillin depletion decreased the CD14 mRNA level and reduced the levels of total and the cell surface CD14. Furthermore, flotillins and mature CD14 were both present in the DRM fraction confirming the possibility of their transbilayer interaction dependent on cholesterol and phosphatidylserine [59, 60]. Our unpublished observations show that CD14 cross-linking in the plasma membrane induces *S*-palmitoylation of flotillin-1 and -2 further underscoring the interdependence of these proteins. On the other hand, the reduction of *Tlr4* expression, minor compared to that of *Cd14*, did not lead to the depletion of TLR4 at the protein level, arguing against a TLR4 deficiency being the cause of the impaired response to LPS.

Flotillins have been indicated to affect gene expression through their influence on the cellular level of sphingosine-1-phosphate (S1P) which in turn inhibits histone deacetylases HDAC1 and HDAC2 [57]. Previous studies of our group indicated that silencing of the gene encoding sphingomyelin synthase 1 (an enzyme converting ceramide into sphingomyelin) in J774 macrophage-like cells also reduced the CD14 mRNA level [61]. Therefore, an influence of flotillin depletion on *Cd14* (and possibly other genes) expression mediated by sphingolipids seems likely. The *Cd14* promoter is regulated (positively and negatively) by the transcription factors Sp1 - Sp3, whose activity, in turn, is regulated by ceramide, reinforcing the possibility of its dysregulation by the disturbances of the flotillins - sphingolipids axis [62–64].

The reduced expression of *Cd14* could be the reason for the paucity of CD14 protein in flotillin-depleted cells, however, disruption in CD14 trafficking was also likely. A line of data indicates that flotillin deficiency can affect the synthesis and anterograde transport of proteins toward the plasma membrane. Thus, a depletion of flotillin-1 induced endoplasmic reticulum stress in HUVEC cells, blocking in turn translation of caveolin-1 mRNA and ultimately TLR3 activation requiring a caveolin-dependent uptake of its ligand, poly-I:C [41]. Of interest, the TLR4-dependent signaling was not affected in those cells lacking endogenous CD14 [41]. In HeLa and HEK293 cells shRNA-mediated silencing of *Flot1* inhibited the trafficking of IGF-1 receptor to the plasma membrane and, as a consequence, the signaling triggered by the receptor [56]. On the other hand, our image analysis of Raw264 cells did not reveal disturbances in CD14 location and abundance in the Golgi apparatus marked by the FAPP1-PH probe. Additionally, a relative enrichment rather than deficiency of CD14 in golgin-97-positive TGN was detected. These data suggest that in contrast to caveolin-1 or IGF-1 receptor, the transport of CD14 from the endoplasmic reticulum toward the plasma membrane was not affected by flotillin depletion.

Somewhat surprisingly, we also did not detect significant changes in the rate of the constitutive endocytosis of CD14 in the flotillin-depleted Raw264 cells. Taking into account the colocalization of flotillin and GPI-anchored proteins in rafts, these proteins were originally considered the most likely cargo of flotillin-dependent clathrin-independent uptake [29, 65]. However, silencing of *Flot1* reduced the endocytosis of CD59 (a GPI-anchored protein) by about 50% only, and also some ultrastructural observations cast doubt on the flotillin involvement in this process. Moreover, flotillin depletion did not inhibit the endocytosis of the T-cell receptor (TCR), a hallmark receptor relying on rafts for signal transduction [36], and our data are in agreement with this line of results. Instead, a crucial role of flotillins in fluid phase marker internalization and also in clathrin-dependent uptake of some proteins has been observed [66, 67], indicating that the cellular functions of flotillins go beyond their contribution to raft dynamics. In agreement, an analysis of the DRM composition in HEK293 cells has revealed that flotillins are found in DRM fractions of higher density resembling “heavy” protein-rich DRM involved in TCR signaling [68]. Also in our analyzes flotillins were present in the DRM, Triton X-100-soluble, and SDS-soluble (cytoskeletal) fractions of Raw264 cells.

The presented results indicated that the depletion of flotillins enhances markedly the recycling of CD14, in line with earlier reports showing that flotillins participate in the recycling of TCR, alpha-amino-3-hydroxy-5-methyl-4-isoxazolepropionic acid (AMPA) receptor, transferrin receptor, E-cadherin, and integrin α-5/β-1. Their flotillin-mediated recycling was required for sustained TCR and AMPA signaling, and the assembly of adherens junctions and focal adhesions [35, 36, 69–71]. Recent proteomic studies identified flotillins in EEA1-positive early endosomes and in lysosomes [38], and the flotillin-mediated recycling of the above-mentioned proteins involved EEA1/Rab5- and Rab11a-positive endosomes. Also, a direct binding of flotillin-2 to Rab11a has been shown [69]. Of note, a flotillin depletion in HeLa and A431 cells up-regulated the recycling of integrin α-5/β-1, resembling the effect on the CD14 recycling observed in our study. It was suggested that the enhanced integrin recycling occurred at the expense of the Rab11a-dependent one, likely in a Rab4-dependent manner [70] Our earlier studies indicated that CD14 recycling is dependent on sorting nexins SNX1, 2, and 6 potentially interacting with the retromer. We found that the constitutive recycling of CD14 in resting cells was crucial for the maintenance of its cell surface level, as a depletion of the SNXs led to a concomitant depletion of the surface CD14. As a result, the TRIF-dependent signaling of TLR4 was inhibited, despite an increase of the TLR4 abundance [22]. Our current microscopic analysis of the CD14 distribution in Raw264 cells indicated that the flotillin depletion led to an enrichment of CD14 in EEA1-positive early endosomes and the TGN compartment involved in some protein recycling routes, while the Rab11-positive ERC compartment seemed not to be involved in the upregulated CD14 recycling. Taken together, the data indicate that while the recycling of plasma membrane proteins via Rab5- and Rab11-positive endosomes requires a flotillin participation, flotillins seem dispensable for the CD14 recycling mediated by SNX1, 2, and 6 and in fact a deficit of flotillins augments this latter CD14 trafficking pathway. Additionally, we observed an enhanced decoration of early endosomes with EEA1 in flotillin-depleted cells. These data suggest that the changes in CD14 recycling possibly followed the changes in the organization of early endosomes, as had been revealed earlier for TCR recycling in flotillin-depleted Jurkat T cells [36].

Paradoxically, despite the enhanced recycling of CD14 in the flotillin-depleted Raw264 cells, its surface level was actually reduced. It seems likely that the enhanced recycling could be a compensatory mechanism striving to maintain the cell surface level of CD14, similarly as it has been suggested for the accelerated recycling of integrin α-5/β-1 in flotillin-depleted cells [70]. The augmented recycling of CD14 relied exclusively on the newly synthesized CD14, while in control cells only a fraction of the recycling CD14 derived from *de novo* synthesis. Based on these data one can infer that after a single round of recycling CD14 was directed from EEA1-positive endosomes to degradation in lysosomes or was removed from the cell surface by shedding. We found that the release of sCD14 can be intensified in the flotillin-depleted cells relative to control ones and was further increased after stimulation with LPS. It has been established that CD14 shedding takes place in resting macrophages and is enhanced after LPS stimulation [72, 73], and that matrix metalloproteinases (MMP)-9 and -12 catalyze this process [74]. A CD14 fragment called presepsin can also be exocytosed after elastase-mediated proteolysis of internalized CD14 [75]. As mentioned above, sCD14 can transfer LPS monomers to the TLR4/MD-2 complex, therefore the efficient shedding of CD14 can explain the low sensitivity of the MyD88-dependent signaling to the deficiency of the membrane-bound CD14 in flotillin-depleted cells.

In conclusion, our data indicate that flotillin depletion leads to a depletion of the total and cell-surface CD14 by down-regulation of *Cd14* expression and stimulating the release of sCD14. Concomitantly, the recycling of newly synthesized CD14 through EEA1-positive early endosomes and TGN is enhanced, possibly aiming to compensate for the deficiency of CD14 on the cell surface. The paucity of surface CD14 weakens the endosomal TRIF-dependent inflammatory signaling of TLR4 after stimulation of cells with LPS.

## Supplementary information

The online version contains supplementary material.

## Acknowledgments

We thank Dr. Jan Fronk (retired), formerly from the Faculty of Biology, University of Warsaw for helpful comments. Confocal imaging was performed at the Laboratory of Imaging Tissue Structure and Function, core facility at the Nencki Institute of Experimental Biology and a part of the infrastructure of the Polish Euro-BioImaging Node. Flow Cytometry was performed at the Laboratory of Cytometry at the Nencki Institute of Experimental Biology.

## Author Contribution

**OVM -** Conceptualization, Investigation, Formal analysis, Writing – review & editing. **AC** - Conceptualization, Methodology, Investigation, Formal analysis, Project administration, Writing - original draft, review & editing. **A H-J** - Investigation, Formal analysis, Writing – review & editing. **NN** - Investigation, Writing – review & editing**. IBA** - Investigation, Writing – review & editing. **KK** - Conceptualization, Supervision, Project administration, Writing – original draft, review & editing.

## Data availability

All data generated or analyzed during this study are included in this article (and its Supplementary Information files) and are available from the corresponding authors on reasonable request.

## Declarations

### Consent for publication

All the authors have read the manuscript and agreed to submit the paper to the Journal.

### Ethics approval

The procedure of obtaining peritoneal macrophages from mice has been reviewed and approved by the Local Animal Ethics Committee (permission No. 394/2017). No research involving humans was conducted and no relevant ethics approval or consent to participate and publish are required.

