## Supplemental data for "Flotillins affect LPS-induced TLR4 signaling by modulating the trafficking and abundance of CD14"

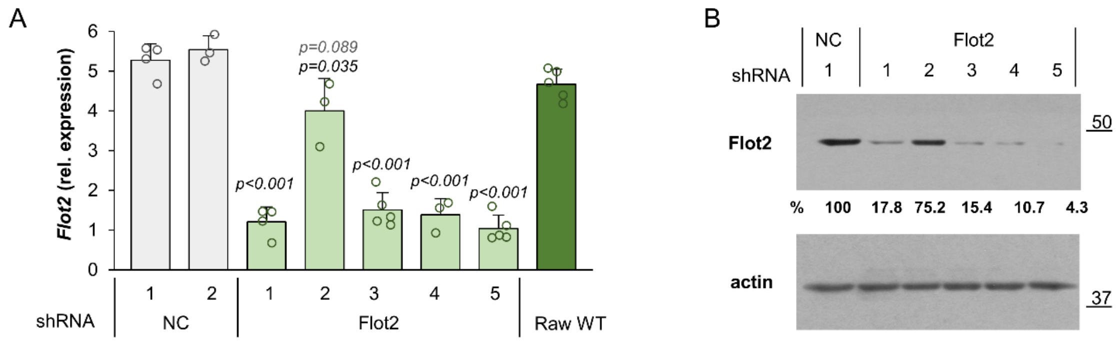


**Supplementary Fig. 1** Knock-down of *Flot2* in Raw264 cells with the application of shRNA. Five commercially available shRNA variants specific against *Flot2* were used individually (No. 1-5) and one control shRNA was applied in two independent approaches (NC1, NC2). Wild type Raw264 cells - Raw WT. (A) RT-qPCR analysis of flotillin-2 mRNA quantified relative to TBP mRNA. Data shown are mean ± SD from at least three experiments. (B) Immunoblotting analysis of flotillin-2 abundance in the indicated clones. Numbers below blots show results of densitometric analysis of flotillin-2 content relative to actin, expressed as the percentage of the value for NC1. Molecular weight markers are shown on the right in kDa. Results of one representative experiment are shown. Significantly different values as indicated by 1-way ANOVA with Scheffe's post hoc test are marked. The *p* value was < 0.001 relative to both NC variants except for Flot2 shRNA variant No. 2 whose upper *p* value relates to NC1 and lower one to NC2

A

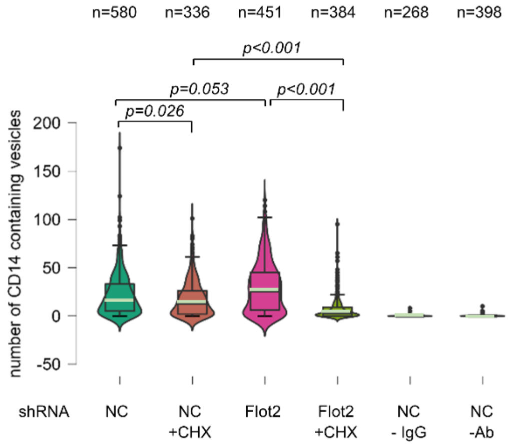


B

| exp. | variant | number of cells | surface CD14 sum of intensities | surface CD14 Intensity / cell | % |
|---|---|---|---|---|---|
| I | NC shRNA | 100 | 4.3203E+10 | 432030821.2 | **100** |
| | Flot2 shRNA | 84 | 2.6775E+10 | 318747168.4 | **73.8** |
| II | NC shRNA | 220 | 5.2199E+10 | 237268251.2 | **100.0** |
| | Flot2 shRNA | 171 | 2.3853E+10 | 139492674.9 | **58.8** |

**Supplementary Fig. 2** CD14 recycling and cell-surface level in flotillin-depleted cells. The recycling was determined in an experiment independent of that shown in Fig. 7 while the data on the level of cell-surface-bound CD14 derive from both these experiments. Cells were transfected with *Flot2*-specific shRNA variant No. 5 (Flot2 shRNA) or control shRNA NC1 (NC shRNA). (A). Quantitation of CD14 recycling. Recycled CD14 was quantitated by a fluorescent antibody-based assay depicted in Fig. 6A. The cells were examined under a fluorescence microscope and vesicles containing fluorescently labeled CD14 representing the recycling CD14 were counted in *n* number of cells. When indicated, the cells were pretreated with 20 µg/mL CHX prior to the assay and CHX was also present during the assay. In control samples, rat isotype IgG2a was used instead of the CD14-specific antibody or the antibody was omitted. Box plots represent median (light green lines) and 25th/75th quartiles of the number of vesicles containing fluorescently labeled CD14 per cell. Significantly different values as indicated by Kruskal-Wallis test followed by Dunn's multiple comparisons post-hoc test are marked. (B) Surface level of CD14 in cells used for the recycling assay. CD14 was visualized on the cell surface with rat anti-CD14 IgG2a followed by chicken anti-rat IgG-Alexa Fluor 647 and quantitated by image analysis using ImageJ

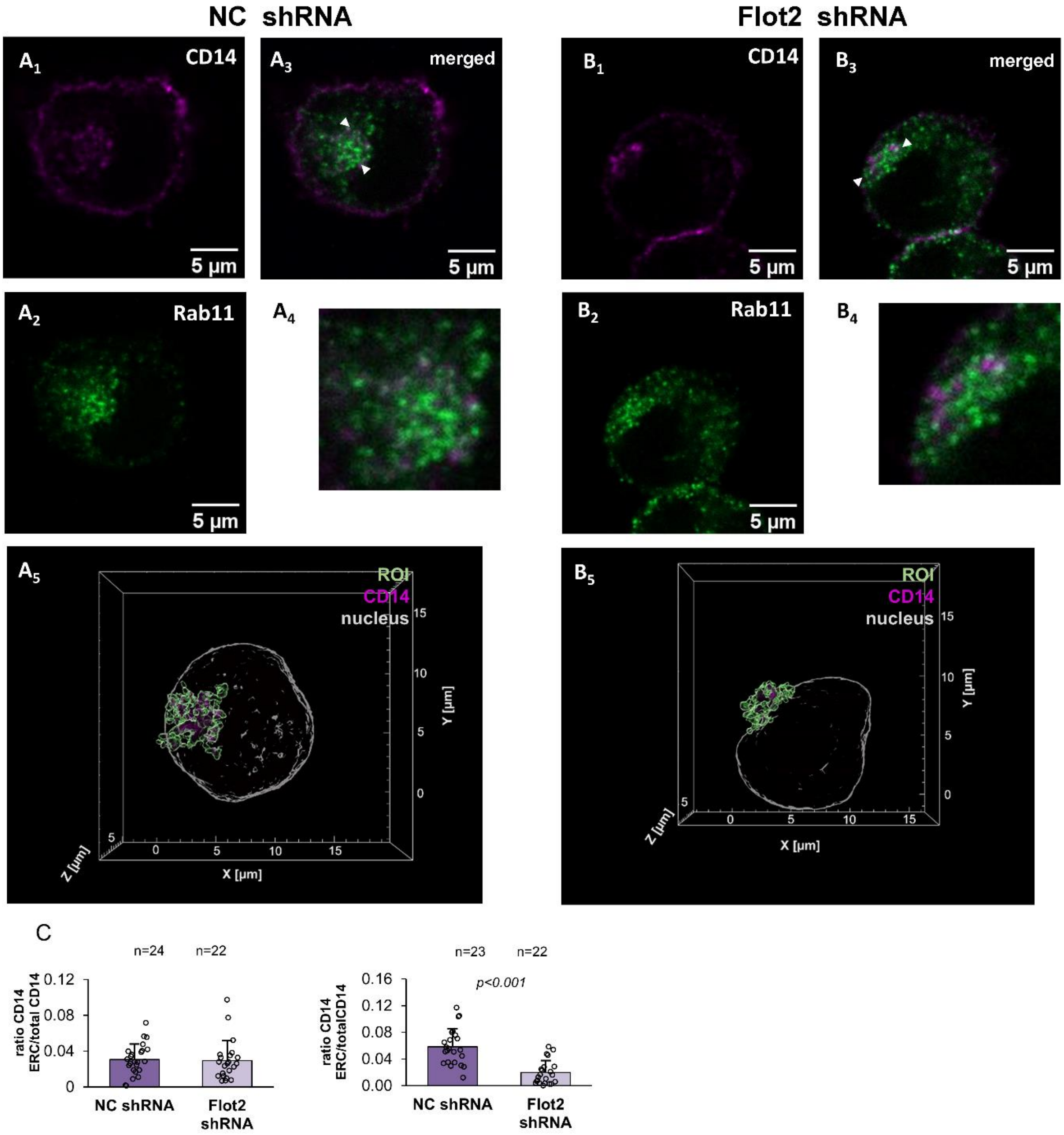


**Supplementary Fig. 3** Enrichment of CD14 in Rab11-positive ERC in flotillin-depleted cells. (A) Raw264 cells transfected with control shRNA, variant NC1. (B) Cells transfected with shRNA specific against *Flot2*, variant No. 5. (A1, B1) Localization of CD14, (A2, B2) Rab11 and (A3, B3) merged images of CD14 and Rab11 localization. z-Stack images of three optical sections taken in the middle of a cell are shown. Colocalized CD14 and Rab11 appear as white spots. (A4, B4) Enlarged images of Rab11-positive ERC indicated by arrowheads in (A3) and (B3). (A5, B5) Reconstructed 3D images of the Rab11-positive ERC delineated in green with CD14 visualized in pink. Contours of the nucleus detected by Hoechst 33342 staining are shown in grey. ROI as these were used for quantitative analysis of CD14 localization in the compartment - see Table 4. (C) Ratio of the fluorescence intensity of CD14 in Rab11-positive ERC calculated as the sum of CD14 voxel intensities to the total CD14 fluorescence detected in z-projections of the respective cell. n, number of cells analyzed. Results of two independent experiments are shown. Significantly different value as indicated by Student's *t*-test is marked

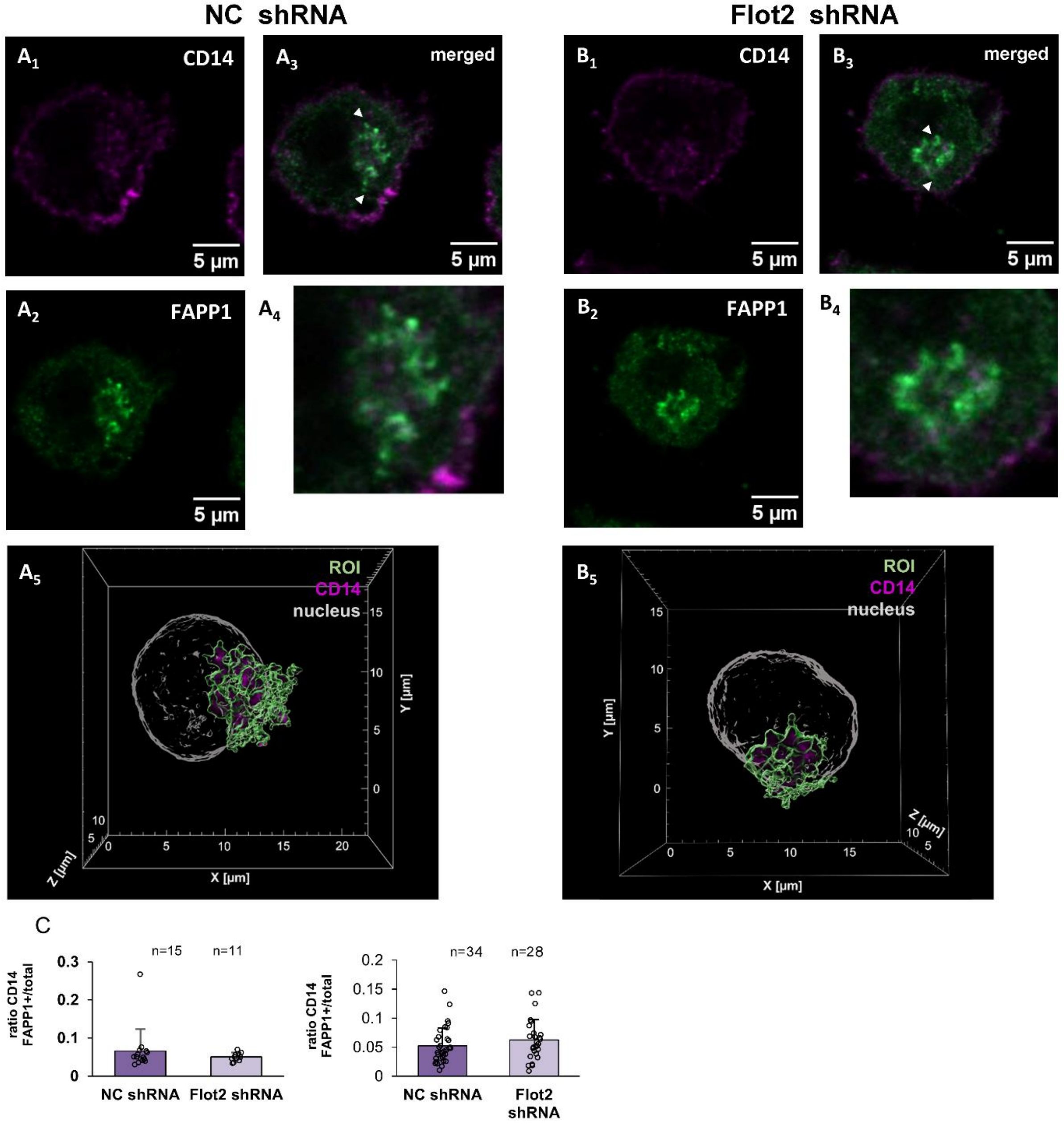


**Supplementary Fig. 4.** Enrichment of CD14 in the Golgi apparatus decorated with FAPP1-PH in flotillin-depleted cells. (A) Raw264 cells transfected with control shRNA, variant NC1. (B) Cells transfected with shRNA specific against *Flot2,* variant No. 5. (A1, B1) Localization of CD14, (A2, B2) FAPP1-PH and (A3, B3) merged images of CD14 and FAPP1-PH localization. z-Stack images of three optical sections taken in the middle of a cell are shown. Colocalized CD14 and FAPP1-PH appear as white spots. (A4, B4) Enlarged images of the Golgi apparatus decorated by FAPP1-PH and indicated by arrowheads in (A3) and (B3). (A5, B5) Reconstructed 3D images of the Golgi apparatus decorated by FAPP1-PH delineated in green with CD14 visualized in pink. Contours of the nucleus detected by Hoechst 33342 staining are shown in grey. ROI as these were used for quantitative analysis of CD14 localization in the compartment - see Table 4. (C) Ratio of the fluorescence intensity of CD14 in the Golgi apparatus calculated as the sum of CD14 voxel intensities to the total CD14 fluorescence detected in z-projections. of the respective cell. n, number of cells analyzed. Results of two independent experiments are shown
